# Massively Parallel Reporter Assay Confirms Regulatory Potential of hQTLs and Reveals Important Variants in Lupus and Other Autoimmune Diseases

**DOI:** 10.1101/2023.08.17.553722

**Authors:** Yao Fu, Jennifer A. Kelly, Jaanam Gopalakrishnan, Richard C. Pelikan, Kandice L. Tessneer, Satish Pasula, Kiely Grundahl, David A. Murphy, Patrick M. Gaffney

## Abstract

**Objective:** To systematically characterize the potential for histone post-translational modifications, i.e., histone quantitative trait loci (hQTLs), expression QTLs (eQTLs), and variants on systemic lupus erythematosus (SLE) and autoimmune (AI) disease risk haplotypes to modulate gene expression in an allele dependent manner.

**Methods:** We designed a massively parallel reporter assay (MPRA) containing ∼32K variants and transfected it into an Epstein-Barr virus transformed B cell line generated from an SLE case.

**Results:** Our study expands our understanding of hQTLs, illustrating that epigenetic QTLs are more likely to contribute to functional mechanisms than eQTLs and other variant types, and a large proportion of hQTLs overlap transcription start sites (TSS) of noncoding RNAs. In addition, we nominate 17 variants (including 11 novel) as putative causal variants for SLE and another 14 for various other AI diseases, prioritizing these variants for future functional studies primary and immortalized B cells.

**Conclusion:** We uncover important insights into the mechanistic relationships between genotype, epigenetics, gene expression, and SLE and AI disease phenotypes.

## Introduction

The non-coding genome is a significant contributor to autoimmune (AI) disease susceptibility and progression (1). Approximately ∼90% of AI disease causal variants are non-coding, with ∼60% mapping to immune cell enhancers and other types of *cis*-regulatory elements (cREs) (2). cREs (e.g., enhancers, promoters, and CTCF-occupied elements (silencers and insulators)) each have unique roles in controlling the expression of nearby genes (3). Histone modifications (e.g., acetylation, methylation, phosphorylation, etc.) also play a pivotal role in gene regulation by influencing chromatin structure and transcription factor (TF) accessibility. Complex interactions between histone modifications and TFs co-localized at cREs influence gene expression (4), and genetic variants located within cREs can alter the binding of TFs, leading to disrupted immune homeostasis.

Our laboratory previously integrated epigenetic and genotypic data from systemic lupus erythematosus (SLE) patient-derived Epstein-Barr virus (EBV)-transformed B cells to assess the degree to which genetic variants in non-coding regions of the genome influence the epigenome (5). We identified variants that affect H3K4me1 and H3K27ac histone modifications, i.e., histone quantitative trait loci (hQTLs). We discovered that H3K4me1 and H3K27ac hQTLs are enriched on AI disease risk haplotypes and disproportionately influenced gene expression variability compared to non-hQTL variants in strong linkage disequilibrium (LD). In an effort to systematically explore the regulatory potential attributed to these hQTL variants, we designed a massively parallel reporter assay (MPRA) to evaluate hQTLs, expression QTLs (eQTLs) shown to interact with hQTLs (5), SLE and AI disease index SNPs, and proxies (r^2^ > 0.80) of the selected variants. MPRA leverages a vector containing a reporter gene (typically green fluorescent protein (GFP)), a promoter, and thousands of barcoded DNA sequences to assess the regulatory activity of the sequence, as well as the functional consequences of genetic variants carried on those sequences (6–8). Determining how non-coding genetic variants alter the activity of regulatory elements and influence gene expression is vital to understanding how such intricate regulatory mechanisms contribute to complex traits and human disease.

## Materials and Methods

### Study population

All experiments were approved by the Institutional Review Board at the Oklahoma Medical Research Foundation (OMRF) prior to initiation. The EBV-transformed B cell line was generated from a non-Hispanic, white, 55 year old, female subject with SLE enrolled in the Lupus Family Registry and Repository (9) and provided by OMRF’s Arthritis and Clinical Immunology Biorepository Core (https://aci-cores.omrf.org/biorepository/). Race and ethnicity were self-reported using a form with fixed categories but confirmed by genetic similarity (via principal components analysis) to other self-reported non-Hispanic white individuals.

### Massively Parallel Reporter Assay

#### Variant selection and oligo generation

A 67,035-oligo library (32,481 variants) (Agilent Technologies) was designed with hQTLs that met a family-wise error rate (FWER)≤0.2 (5), hQTL proxies (r^2^≥0.8), published SLE or AI disease index SNPs (p≤5E-08), eQTLs found to interact with hQTLs (5), eQTL proxies, and additional SNPs located on AI haplotypes that included a hQTL. Location controls and random controls were also included. Location controls met the following criteria: 1) minor allele frequency (MAF≥5%); 2) location within 150-1000bp of a hQTL SNP; 3) low LD with hQTL SNP (r^2^≤0.25); and 4) no significant eQTL signal in public eQTL databases. Random control SNPs were selected randomly across the genome and matched the MAF distribution of the hQTL SNPs. Oligos were generated using 150bp of hg19 genomic sequence flanking the reference and alternate alleles of each selected variant (74bp 5’ and 75bp 3’ of the allele of interest) with 15bp adapters added to each end (5′ ACTGGCCGCTTGACG [150bp oligo] CACTGCGGCTCCTGC 3′) **(Supplementary Table 1 (Oligo sequences))**.

#### Plasmid Library Construction

MPRA was performed as described previously (6, 10) with minor modifications. First, oligo-barcode libraries were constructed by 28X parallel PCR reactions to add 20bp random barcodes to the synthesized 180bp oligos. MpraΔorf libraries were assembled using Gibson Assemble Master Mix (NEB E2611L). The GFP amplicon containing a minimal promoter, GFP open reading frame, and partial 3’UTR was amplified from the pGL4.23:minP GFP plasmid and inserted into purified mpraΔorf plasmids by Gibson Assembly. Constructed mpra:gfp libraries were transformed into NEB 10-β *E. coli* (NEBC3020K) by electroporation and expanded in 5L LB media supplemented with 100µg/mL carbenicillin at 37°C, shaking for 16 hours. The mpra:gfp plasmid libraries were purified using the Qiagen Plasmid Plus Giga Kit.

#### MPRA Library Transfections

EBV B cells were cultured in RPMI medium supplemented with 15% FBS, 100U/mL penicillin, 100µg/mL streptomycin, and 2mM L-glutamine at 37°C, 5% CO_2_. Cells were seeded at 5×10^5^ cells-per-mL 36 hours before transfection. Cells were collected and split into six transfections with 100 million cells and 100µg mpra:gfp plasmid library per replicate. Transfection was performed with the Neon transfection system in 100µL tips containing 10 million cells per tip with 3 pulses of 1200V and 20ms each. After transfection, cells were cultured with RPMI supplemented with 15% FBS and without antibiotics for 24 hours. Cells were then collected and lysed in RLT buffer (Qiagen Midi RNeasy 75144) by passing through 18-gauge needles. Cell lysate was stored in −80°C until RNA purification.

#### MPRA Library Complexity Validation

The fragment containing the oligo-barcode combination was amplified from mpraΔorf plasmids and attached to Illumina sequencing adapters with the Illumina TruSeq Universal Adapter and unique P7 index primers. Libraries were sequenced using 2×150 PE reads on the Illumina Novaseq platform.

#### Sequencing Library Preparations

Total cell lysis was thawed on ice and lysed again by passing 5-10 times through 18-gauge needles. GFP mRNA extraction, pull down, and cDNA synthesis was performed as previously described (6, 10). Plasmid libraries and cDNA samples were amplified, and Illumina sequencing adaptors were added using the Illumina TruSeq Universal Adapter and TruSeq_Index primer (NEB E7335S). Libraries were sequenced using the Illumina NovaSeq PE150 targeting 400 million reads per sample.

#### Promoter Capture HiC

Leukoreduction chambers were obtained from the Oklahoma Blood Institute. Primary B cells were isolated with negative magnetic bead selection (StemCell 19054). Cells were seeded at 3×10^6^ cells per mL in RPMI medium supplemented with 10% FBS, 100U/mL penicillin, 100µg/mL streptomycin, and 2mM L-glutamine. After one hour incubation at 37°C, 5% CO_2_, cells were treated with 5µg/mL R837 (TLR7 agonist), 1µg/mL CD40 ligand, and 3µg/mL IgG/M for 48 hours to induce SLE-like inflammation responses. HiC libraries were generated with a pool of 5 million B cells from 10 donors following the Hi-C 3.0 protocol (11). Capture enrichment was performed using the Arima Human Promoter Panel kit (Arima A510008 and A302010) following the manufacturer’s instructions. Libraries were sequenced using the Illumina NovaSeq PE150 targeting 200 million reads per sample. Data were processed following the Arima Genomics pipeline (https://github.com/ArimaGenomics/CHiC) and interactions were viewed on the WashU Epigenome Browser (http://epigenomegateway.wustl.edu/browser/).

### Data Analysis

#### Oligo-barcode associations and barcode counting

Paired oligo-barcode associations were determined in the four sequenced mpraΔorf plasmid control libraries using analysis scripts (e.g. MPRAmatch.wdl) and the pipeline designed by Dr. Ryan Tewhey (https://github.com/tewhey-lab/MPRA_oligo_barcode_pipeline) (6). Pairs with alignment score error rates of greater than 5% were discarded. Only barcodes that uniquely mapped to one oligo were used for downstream analysis. Oligo-barcode pairs from the four libraries were then merged together for a total of 66,949 (99.87%) oligos captured and 145M total oligo-barcode pairs in the initial oligo-barcode pools. Barcode read counting for each oligo-barcode pair was determined in each of the four plasmid control replicates and six transfected EBV B cell line replicates using the MPRAcount.wdl script. Reads were totaled across all barcodes associated with each oligo.

#### Expression modulating variant (EmVar) and allelic effect (AE) variant identification

Oligo counts were normalized by DESeq2 (12) and modeled as a negative binomial distribution to obtain estimates of variance in oligo counts across all samples. EmVars – defined as variants where one or more alleles altered GFP expression (emAlleles) – were determined for the plasmid controls and library replicates and tested for significant expression differences using a Wald’s test (6). A fold change (FC) difference of 1.5 between the plasmid controls and EBV library replicates and an FDR<0.05 were required for significance. EmVars were then assessed for allele-specific transactivation potential (allelic effects, AE) by comparing log_2_ ratios of the reference versus alternate alleles using a Student’s t-test (6). A FC=1.25 between the two alleles and an FDR<0.05 were required for significance. For the 320 multiallelic variants, the reference allele was compared to each alternate allele separately.

#### Transcription factor (TF) motif enrichment analysis

The findMotifs.pl program within HOMER (13) was utilized to evaluate emVars and hQTL emVars for known TF motif enrichment. To evaluate the emVars, emVar oligo FASTA sequences were used as target sequences and non-emVar oligo FASTA sequences were used as background sequences; for the hQTL emVar analysis, target FASTA sequences for hQTL emVar oligos were compared to background FASTA sequences for non-hQTL emVar oligos.

#### Annotations

Prior to downstream analysis, variant positions were converted to hg38 using the UCSC LiftOver tool (https://genome.ucsc.edu/cgi-bin/hgLiftOver). Human ENCODE CRE information (encodeCcreCombined.bb) (3) was downloaded from the UCSC genome browser (http://hgdownload.soe.ucsc.edu), converted to a bed file using the bigBedToBed tool, and evaluated for overlaps with MPRA variant locations for annotation. SLE and AI index SNP identification was initially obtained from the NHGRI-EBI GWAS catalog (14) in 2018; annotations were later updated using the NHGRI-EBI GWAS catalog v1.0.2 download on 03-27-2023. Published index SNPs (p<5E-8) from the following AI diseases were identified: ankylosing spondylitis (AS), autoimmune thyroid disease (Graves’ disease or Hashimoto’s thyroiditis), celiac disease, Crohn’s disease (CD), Kawasaki disease (KD), multiple sclerosis (MS), myasthenia gravis (MG), myositis, primary biliary cirrhosis (PBC), psoriasis, rheumatoid arthritis (RA), sarcoidosis, Sjögren’s disease (SjD), systemic sclerosis (scleroderma, SS), type 1 diabetes (T1D), ulcerative colitis (UC), and vitiligo. Mapped and nearest gene annotations were obtained from Ensembl’s variant effect predictor tool (https://useast.ensembl.org/Homo_sapiens/Tools/VEP; release 109, February 2023) (15).

## Results

### hQTLs demonstrate strong regulatory activity, are enriched for interferon regulatory factor TFs, and are located in regions that commonly overlap non-coding RNAs

A total of 66,865 (99.87%) oligos were recovered from the plasmid control and EBV B cell line libraries with sufficient oligos having >10 barcodes per oligo (98.9% and 91.3% in the plasmid controls and EBV B cell replicates, respectively) and >20 mean reads count per oligo (98% and 96.2% in the plasmid controls and transfected EBV B cell replicates, respectively) (**Supplementary** Figures 1A-D). The experimental replicates of each library produced strong reproducibility of the normalized read counts, and the EBV B cell samples produced more normalized reads per oligo, on average, than the plasmid controls (**Supplementary** Figures 1E **& F)**. Oligos with <20 associated barcodes (n=1,475) and <20 average reads in the plasmid controls (n=569) were removed from the downstream analysis, resulting in a total of 64,821 oligos with an average coverage of 1,271 barcodes/oligo and 31,666 total variants for analysis.

Our post-QC dataset included 4,039 hQTL variants; 6,810 hQTL proxies; 163 SLE index SNPs (p ≤5E-8); 802 AI index SNPs (p≤5E-8); 1,198 SLE and AI index SNP proxies (r^2^>0.8); 2,001 eQTLs; 11,793 eQTL proxies; 1,526 proxies of a hQTL located on an AI haplotype (“haplotype SNPs”); 1,968 location controls; 975 random controls; and 425 additional AI index SNPs with p≤1E-6 (“suggestive AI SNPs”). We found that 4,780 (15.1%) of the tested variants were emVars, variants for which one or more alleles (n=7,911, “emAlleles”) demonstrated regulatory activity of the GFP reporter (**Figure 1A** and **Supplementary Table 2).** The vast majority of emAlleles (n=7,898; 99.8%) exhibited higher regulatory activity in the EBV B libraries compared to controls; this was not unexpected, however, since we used a construct with a low basal activity promoter making it easier to detect inducible effects (6) **(Figure 1B)**. When evaluating the different types of variants, hQTLs produced the highest proportion of emVars (24%, n = 976), followed by published SLE index SNPs (20%, n=32) (**Figure 1C)**. While there was no significant difference in emVar proportions between hQTLs and SLE risk SNPs, the proportion of hQTL emVars was significantly higher than every other variant type tested including AI disease index SNPs (15%, n= 25, χ^2^=28.02, p<0.00001), and eQTLs (14%, n=286, χ^2^=78.89, p<0.00001). hQTLs also displayed the strongest effects on gene expression compared to other tested variant types, with mean and median EBV B/control FC=3.06 (log_2_(FC)=1.61)) and 2.60 (log_2_(FC)=1.36), respectively, demonstrating that many hQTLs are capable of inducing significantly stronger gene expression activity than eQTLs (mean and median FC=2.60 and 2.27, respectively, t_mean_=5.43, p<0.0001) and other variants in strong LD with them **(Figure 1D)**.

**Figure 1.**
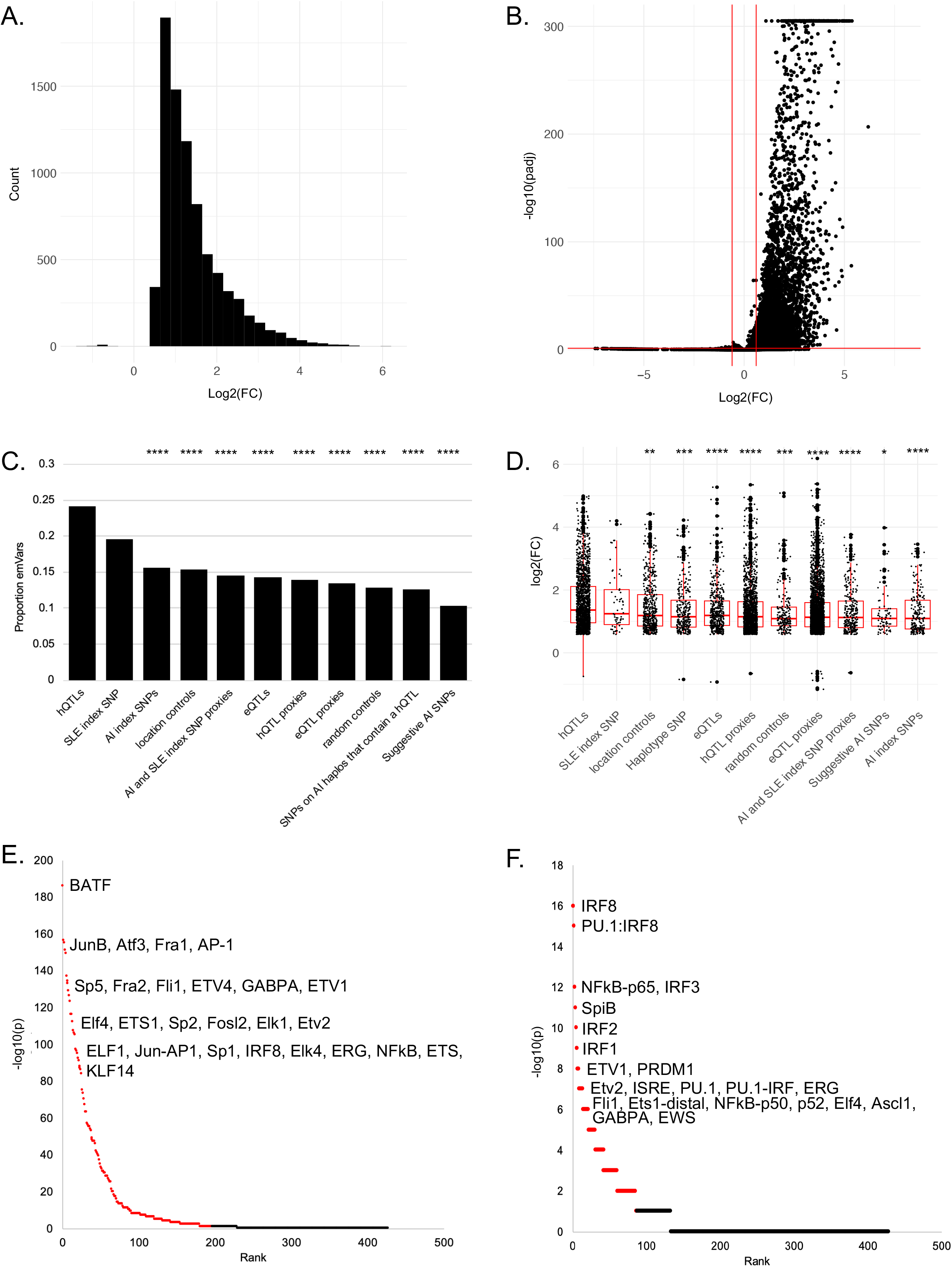
Properties of emVars in the MPRA. **A.** Histogram distribution of emVar regulatory (log_2_(FC)) in six EBV B cell technical replicates compared to four plasmid controls. Positive values represent increased regulatory activity and negative values represent decreased activity in EBV B cells relative to plasmid controls. Oligo count is plotted on the y-axis. **B.** Volcano plot of emVar effect sizes (−log_10_(p_adj_) from DESeq2) in EBV B cells relative to controls. Horizontal red line represents p_adj_≤0.05; vertical red lines (log_2_(FC)±0.58) represent a 1.5 FC difference between the EBV B replicates and plasmid controls. **C.** Proportion of emVars within each variant type. Significant differences between the proportion of emVars within hQTLs and the other variant types are shown: ****chi-square p<0.0001. **D.** Box plots of emVar effect sizes (log_2_(FC)) for each variant type. X-axis is sorted in descending order by mean log_2_(FC) of the variant types. Significant differences in the means of hQTL effect sizes compared to the other variant types are shown: *t-test p<0.05; **<0.01, ***<0.001, ****<0.0001. **E & F.** Significant TFs enriched in all emVars (E) and hQTL emVars (F). TF rank and HOMER-log_10_(p) are plotted. FDR≤0.05 effects are highlighted in red. Top TFs are indicated.

When emVars were tested for enrichment of TF motifs compared to the 26,886 variants that did not exhibit effects on expression, an enrichment of 196 TFs in the emVars was observed with most being members of the C2H2-zinc finger (ZF; n=32), basic leucine zipper (bZIP; n=27), basic helix-loop-helix (bHLH; n=25), or homeobox (n=25) and erythroblast transformation specific (ETS; n=25) families. The most strongly enriched TFs within the emVars, however, were part of the bZIP and ETS families (**Figure 1E** and **Supplementary Table 3).** The strongest bZIP family TFs included *BATF* (p=1E-186; 11.25% of emVar sequences included the motif vs 3.16% in non-emVars)*, JunB* (p=1E-156; 9.28% vs 2.55%)*, Atf3* (p=1E-155; 10.52% vs. 3.23%), and others. Strong ETS family TFs included *Fli1* (p=1E-133; 13.85% in emVars vs 5.68% in non-emVars), *ETV4* (p=1E-129; 13.24% vs 5.37%), *GABPA* (p=1E-126; 11.57% vs 4.37%), and others (**Figure 1E** and **Supplementary Table 3)**. When we narrowed the analysis further to identify TFs that affect gene expression within hQTLs emVars compared to non-hQTL emVars, we observed enrichment of 87 TFs, with strongest enrichment among TF members of the interferon regulatory factor (IRF) family (e.g., *IRF8, IRF3, IRF2, IRF1, ISRE,* etc.), indicating that these hQTLs are potentially important in the regulation of interferon signaling responses in B cells (**Figure 1F** and **Supplementary Table 4)**.

A significantly higher proportion of emVars were located in ENCODE cREs than non-emVars (84% vs 80%; χ^2^=64.9; p<0.00001) (**Supplementary** Figure 2A). EmVars were most commonly found within ENCODE’s enhancer like signatures (ELS) characterized by high DNase and H3K27ac activity and low H3K4m3 activity (3), as well as more frequently located >2kB of an annotated TSS (distal ELS, dELS, 61.3%) than within 2kB of an annotated TSS (promoter ELS, pELS, 13%), indicating that most emVars likely overlap TSS of noncoding RNAs (3). We also found that 91% of hQTL emVars were located in a cRE, which was not unexpected since they were originally identified as H3K27ac and H3K4me1 hQTLs (5), and, similar to emVars as a group, likely overlap TSS of noncoding RNAs (70% in a dELS) **(Supplementary** Figures 2B**&C)**. While a high proportion of SLE index emVars were also found in cREs (90%, n=57), only 50% were located in dELS, and had the highest proportion of variants located in DNase-H3K4me3 (6%) and CTCF-only (6%) cREs compared to the other variants tested, suggesting that a proportion of these variants likely exhibit different regulatory functions than hQTLs and the other types of variants tested. (**Supplementary** Figures 2D-G).

### EmVars with strong allelic effects are dominated by hQTLs

We evaluated the 4,765 (99.7%) emVars that had both alleles represented for significant differential regulatory effects between the reference and alternate alleles (allelic effects, AE variants). A total of 567 emVars (11.9%; 572 alleles due to multiallelic variants) were AE variants with 28.5% (n=162) boasting strong allele-specific differences in expression (FC >2; log_2_(FC) >1 or <-1); of these, 45% (n=73) were hQTLs, 23% (n=38) were eQTL proxies, and 24% (n=15) were hQTL proxies (**Figures 2A & B** and **Supplementary Table 5**). A total of 299 variants (302 alleles due to multiallelic variants) displayed significant increases in expression with the reference allele and 270 with the alternate allele **(Figure 2A)**. When looking at the different types of variants evaluated, hQTLs and published SLE index SNPs produced the highest proportion of AE variants (21% (201/971) and 20% (7/35), respectively) (**Figure 2C).** When comparing effects driven by the alternate or reference allele, the alternate allele more often demonstrated higher expression (median log_2_(FC) >0) for AI and SLE index SNPs and eQTLs, while the reference allele more often demonstrated significantly higher expression (median log_2_(FC)<0) in the other variant types (**Figure 2D).**

**Figure 2.**
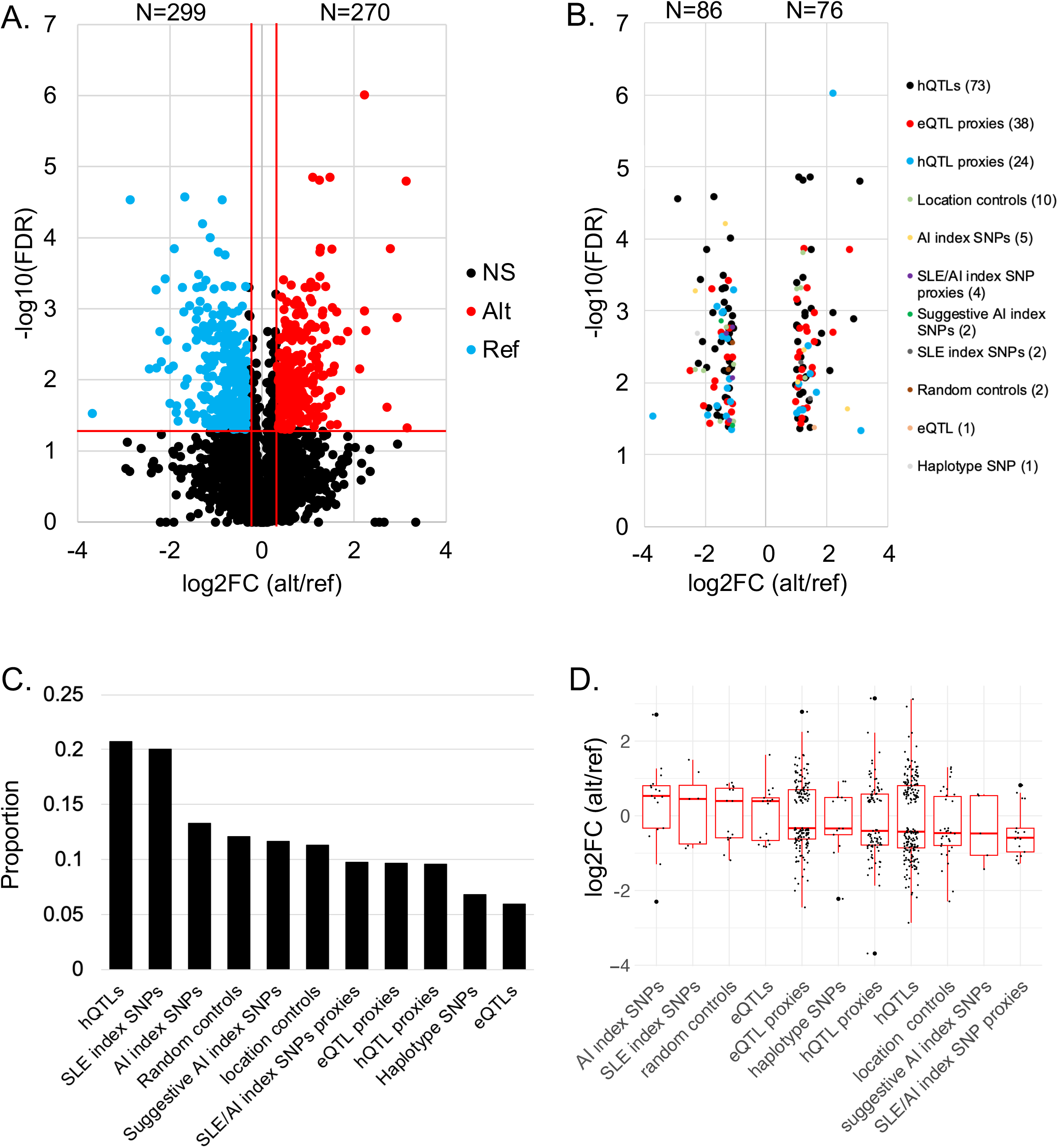
Strong AE variants are dominated by hQTLs. **A.** Volcano plot of AE effect sizes (log_2_(FC)) relative to the alternate/reference alleles. Horizontal red line represents q£0.05; vertical red lines (log_2_(FC)=±0.32) represent a 1.25 FC difference between the alternate/reference alleles. Red dots indicate alternate alleles that demonstrate significantly higher expression than the reference allele; blue dots indicate reference alleles that have significantly higher expression. **B.** AE variants (n=162) with effect sizes >2.0, colored by variant type, demonstrating the high number of hQTLs (45%, black dots) with strong effects. The log_2_(FC) of alternate/reference allele is plotted on the x-axis; y-axis is the −log_10_(FDR). **C.** Proportions of AEs within each type. **D.** The log_2_(FC) of alternate/reference allele effect size plotted by variant type, sorted by descending median log_2_FC alternate/reference.

### AE emVars identify candidate causal variants for SLE and AI disease risk haplotypes

We next focused on effects located on SLE and AI risk haplotypes to identify putative causal variants for disease. In addition to the selected 163 SLE index SNPs that passed QC, 2,567 tested variants were located on SLE risk haplotypes (D’>0.8 to an SLE index SNP). A total of 381 (14%) variants were emVars (35 SLE index SNPs and 346 SLE haplotype SNPs) located on 50 SLE risk haplotypes; 208 of which were within the HLA region (**Figure 3A**). A total of 35 (9%) emVars (17 outside of the HLA region) also demonstrated significant differential expression activity between the two alleles and are, thus, putative causal variant candidates for SLE risk haplotypes. When evaluating the 17 non-HLA AE variants, six are SLE index SNPs **(Table 1**, **Figure 3B, Supplementary** Figures 3A-F **& 4A-F**, and **Supplementary Table 5)** and 11 are novel SLE variant candidates (**Table 1**, **Figures 3B, 4 & 5, Supplementary** Figures 3G-K **& 4G-K** and **Supplementary Table 5)**; 16 have a RegulomeDb score of 1, providing additional support for their high likelihood of being functional (**Supplementary Table 6).** We also identified 14 putative causal variants for index variants for RA, T1D, MS, UC, CD, KD, PBC, GD and vitiligo **(Table 1, Supplementary Table 5,** and **Supplementary** Figure 5). We focus discussion on several novel SLE candidate causal variants identified below.

**Figure 3.**
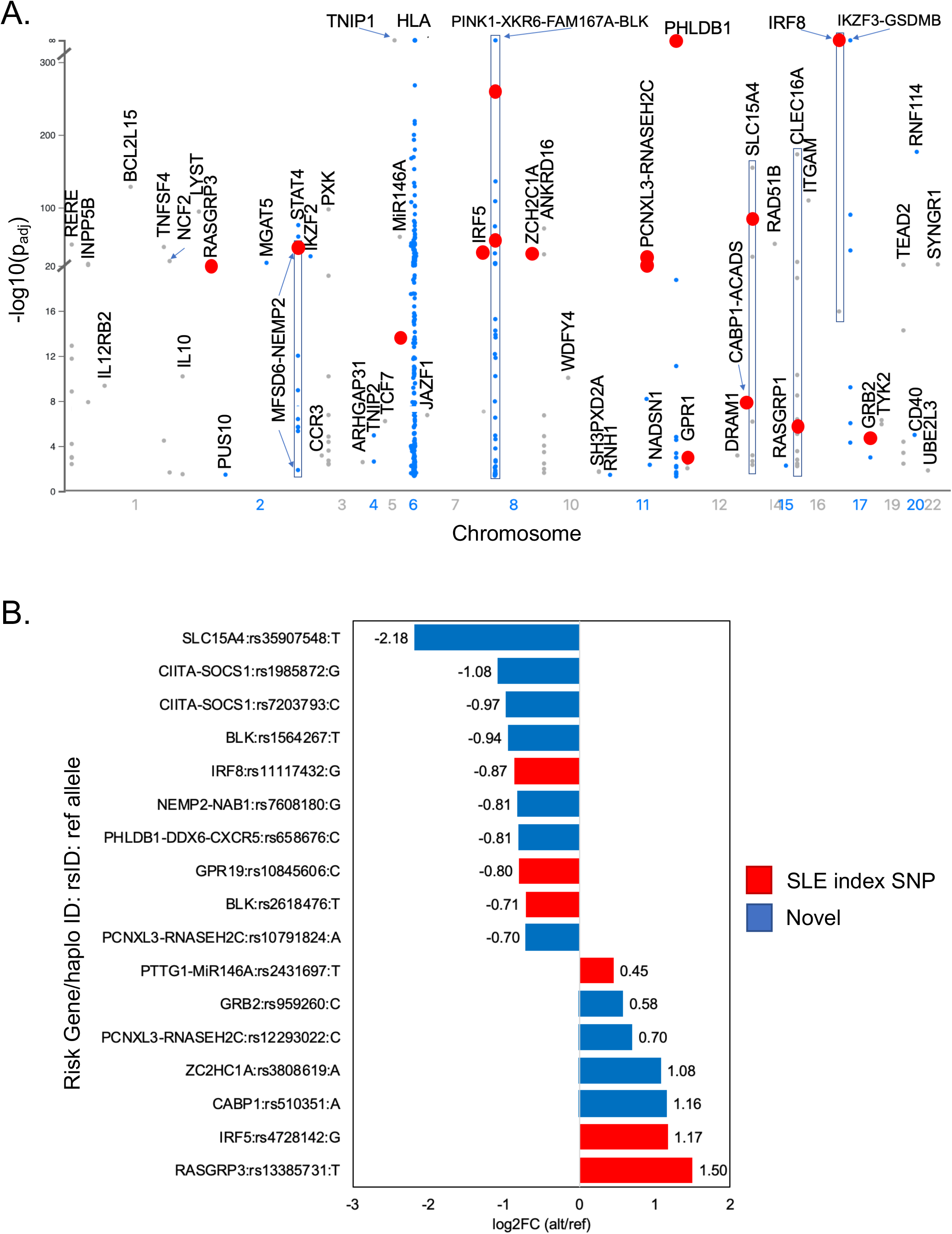
EmVar and AE variants on SLE risk haplotypes. **A.** Manhattan plot of significant emVars (-log_10_(p_adj_) as determined by DESeq2) plotted on SLE risk haplotypes. Only the allele with the highest regulatory activity is plotted for each variant. SLE risk gene/haplotype is indicated. Red dots represent emVars with allelic effects. **B.** Effect sizes of variants that demonstrate significant AEs on SLE risk haplotypes. The risk gene/ haplotype and reference allele are provided for each variant. Positive effects indicate the alternate allele significantly increases expression over the reference allele; negative effects indicate the reference allele significantly increases expression over the alternate allele. Candidate causal SLE index SNPs are indicated in red. Novel candidate causal variants on SLE haplotypes are indicated in blue.

**Table 1.**
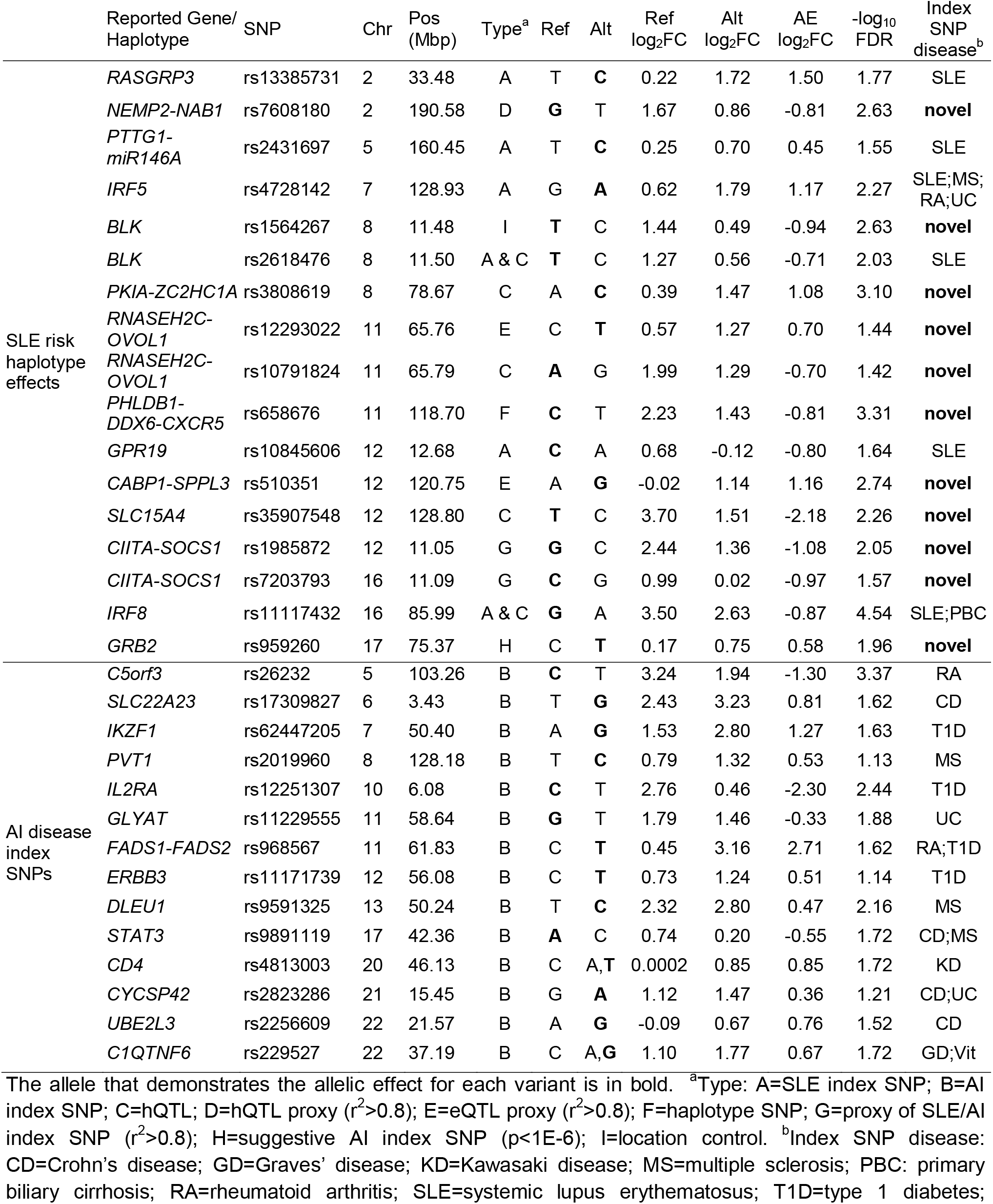
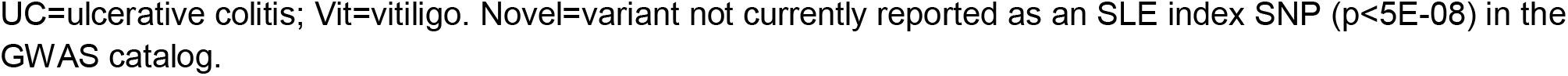
Non-HLA candidate causal variants for SLE and AI disease risk genes/ haplotypes.

|  | Reported Gene/<br>Haplotype | SNP | Chr | Pos<br>(Mbp) | Type<sup>a</sup> | Ref | Alt | Ref<br>log<sub>2</sub>FC | Alt<br>log<sub>2</sub>FC | AE<br>log<sub>2</sub>FC | -log<sub>10</sub><br>FDR | Index<br>SNP<br>disease<sup>b</sup> |
| --- | --- | --- | --- | --- | --- | --- | --- | --- | --- | --- | --- | --- |
| SLE risk<br>haplotype<br>effects | *RASGRP3* | rs13385731 | 2 | 33.48 | A | T | **C** | 0.22 | 1.72 | 1.50 | 1.77 | SLE |
|  | *NEMP2-NAB1* | rs7608180 | 2 | 190.58 | D | **G** | T | 1.67 | 0.86 | -0.81 | 2.63 | novel |
|  | *PTTG1-miR146A* | rs2431697 | 5 | 160.45 | A | T | **C** | 0.25 | 0.70 | 0.45 | 1.55 | SLE |
|  | *IRF5* | rs4728142 | 7 | 128.93 | A | G | **A** | 0.62 | 1.79 | 1.17 | 2.27 | SLE;MS;<br>RA;UC |
|  | *BLK* | rs1564267 | 8 | 11.48 | I | **T** | C | 1.44 | 0.49 | -0.94 | 2.63 | novel |
|  | *BLK* | rs2618476 | 8 | 11.50 | A & C | **T** | C | 1.27 | 0.56 | -0.71 | 2.03 | SLE |
|  | *PKIA-ZC2HC1A* | rs3808619 | 8 | 78.67 | C | A | **C** | 0.39 | 1.47 | 1.08 | 3.10 | novel |
|  | *RNASEH2C-OVOL1* | rs12293022 | 11 | 65.76 | E | C | **T** | 0.57 | 1.27 | 0.70 | 1.44 | novel |
|  | *RNASEH2C-OVOL1* | rs10791824 | 11 | 65.79 | C | **A** | G | 1.99 | 1.29 | -0.70 | 1.42 | novel |
|  | *PHLDB1-DDX6-CXCR5* | rs658676 | 11 | 118.70 | F | **C** | T | 2.23 | 1.43 | -0.81 | 3.31 | novel |
|  | *GPR19* | rs10845606 | 12 | 12.68 | A | **C** | A | 0.68 | -0.12 | -0.80 | 1.64 | SLE |
|  | *CABP1-SPPL3* | rs510351 | 12 | 120.75 | E | A | **G** | -0.02 | 1.14 | 1.16 | 2.74 | novel |
|  | *SLC15A4* | rs35907548 | 12 | 128.80 | C | **T** | C | 3.70 | 1.51 | -2.18 | 2.26 | novel |
|  | *CIITA-SOCS1* | rs1985872 | 12 | 11.05 | G | **G** | C | 2.44 | 1.36 | -1.08 | 2.05 | novel |
|  | *CIITA-SOCS1* | rs7203793 | 16 | 11.09 | G | **C** | G | 0.99 | 0.02 | -0.97 | 1.57 | novel |
|  | *IRF8* | rs11117432 | 16 | 85.99 | A & C | **G** | A | 3.50 | 2.63 | -0.87 | 4.54 | SLE;PBC |
|  | *GRB2* | rs959260 | 17 | 75.37 | H | C | **T** | 0.17 | 0.75 | 0.58 | 1.96 | novel |
| AI disease<br>index<br>SNPs | *C5orf3* | rs26232 | 5 | 103.26 | B | **C** | T | 3.24 | 1.94 | -1.30 | 3.37 | RA |
|  | *SLC22A23* | rs17309827 | 6 | 3.43 | B | T | **G** | 2.43 | 3.23 | 0.81 | 1.62 | CD |
|  | *IKZF1* | rs62447205 | 7 | 50.40 | B | A | **G** | 1.53 | 2.80 | 1.27 | 1.63 | T1D |
|  | *PVT1* | rs2019960 | 8 | 128.18 | B | T | **C** | 0.79 | 1.32 | 0.53 | 1.13 | MS |
|  | *IL2RA* | rs12251307 | 10 | 6.08 | B | **C** | T | 2.76 | 0.46 | -2.30 | 2.44 | T1D |
|  | *GLYAT* | rs11229555 | 11 | 58.64 | B | **G** | T | 1.79 | 1.46 | -0.33 | 1.88 | UC |
|  | *FADS1-FADS2* | rs968567 | 11 | 61.83 | B | C | **T** | 0.45 | 3.16 | 2.71 | 1.62 | RA;T1D |
|  | *ERBB3* | rs11171739 | 12 | 56.08 | B | C | **T** | 0.73 | 1.24 | 0.51 | 1.14 | T1D |
|  | *DLEU1* | rs9591325 | 13 | 50.24 | B | T | **C** | 2.32 | 2.80 | 0.47 | 2.16 | MS |
|  | *STAT3* | rs9891119 | 17 | 42.36 | B | **A** | C | 0.74 | 0.20 | -0.55 | 1.72 | CD;MS |
|  | *CD4* | rs4813003 | 20 | 46.13 | B | C | A,**T** | 0.0002 | 0.85 | 0.85 | 1.72 | KD |
|  | *CYCSP42* | rs2823286 | 21 | 15.45 | B | G | **A** | 1.12 | 1.47 | 0.36 | 1.21 | CD;UC |
|  | *UBE2L3* | rs2256609 | 22 | 21.57 | B | A | **G** | -0.09 | 0.67 | 0.76 | 1.52 | CD |
|  | *C1QTNF6* | rs229527 | 22 | 37.19 | B | C | A,**G** | 1.10 | 1.77 | 0.67 | 1.72 | GD;Vit |

The allele that demonstrates the allelic effect for each variant is in bold. <sup>a</sup>Type: A=SLE index SNP; B=AI index SNP; C=hQTL; D=hQTL proxy ( $r^2>0.8$ ); E=eQTL proxy ( $r^2>0.8$ ); F=haplotype SNP; G=proxy of SLE/AI index SNP ( $r^2>0.8$ ); H=suggestive AI index SNP ( $p<1E-6$ ); I=location control. <sup>b</sup>Index SNP disease: CD=Crohn's disease; GD=Graves' disease; KD=Kawasaki disease; MS=multiple sclerosis; PBC: primary biliary cirrhosis; RA=rheumatoid arthritis; SLE=systemic lupus erythematosus; T1D=type 1 diabetes;
UC=ulcerative colitis; Vit=vitiligo. Novel=variant not currently reported as an SLE index SNP ( $p < 5E-08$ ) in the GWAS catalog.

There is currently one reported SLE index SNP in the region between *NEMP2* and *NAB1* on chromosome 2: rs9630991 (16). We evaluated this variant along with 153 other variants on the haplotype and, while multiple variants were shown to be emVars (**Figure 4A** and **Supplementary Table 2)**, only rs7608180, a variant 11,264bp downstream of rs9630991 (D’=0.95, r^2^=0.35), produced an AE with the reference G allele (FC=0.57, FDR q=0.002) (**Figures 3B & 4B,** and **Supplementary Table 5).** Our promoter capture HiC data collected in primary B cells demonstrates that, while the region containing rs7608180 lies between *NEMP2* and *NAB1,* it interacts with the promoter of the upstream major facilitator superfamily domain containing 6 (*MFSD6*) gene, a gene predicted to enable MHC class I protein binding and receptor activity (17), prioritizing *MFSD6* as a possible SLE risk locus in the region **(Figure 4C).**

**Figure 4.**
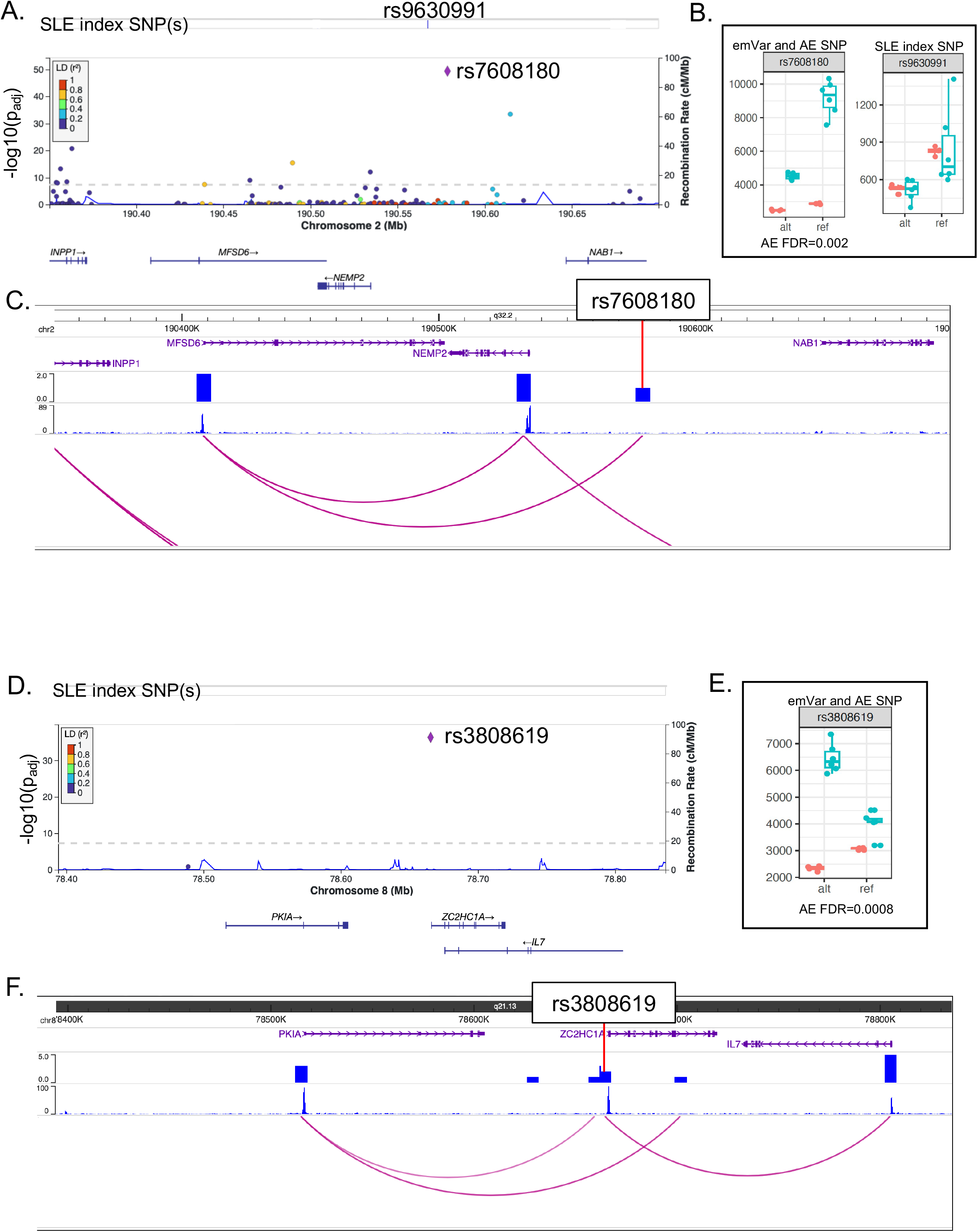
Novel candidate variant identification on the *NEMP2*-*NAB1* and *PKIA*-*ZC2HC1A* SLE risk haplotypes. **A & D.** LocusZoom plots demonstrating emVar effects on the haplotypes of (A) *NEMP2*-*NAB1* and (D) *PKIA*-*ZC2HC1A.* Evaluated index SNPs are indicated. Variants evaluated, their genomic location, and genes in the region are presented on the x-axis. The novel AE variant is represented as a purple diamond. LD values between each variant and the AE variant are colored based on their r^2^ value (see LD key). **B & E.** Box plots of the counts for each allele (alt and ref) in the EBV B (blue) and plasmid control (orange) replicates. FDR q value is indicated. **C & F.** Screenshot from the WashU Epigenome Browser for each haplotype region. Gene positions are provided on the top, followed by density HiC bed tracks, HiC bigwig tracks, and the interactions between gene promoters and the HiC data. The AE variant position is provided.

Association between SLE and rs4739134 has been reported between the region of *PKIA* and *ZC2HC1A* on chromosome 8 (18). We, unfortunately, did not include this variant in our study; however, we did evaluate an hQTL variant (rs3808619) that is strongly correlated with rs4739134 (D’=0.99, r^2^=0.98). rs3808619, located ∼600bp downstream of *ZC2HC1A,* produced a strong emVar effect **(Figure 4D** and **Supplementary Table 2)** with the alternate C allele producing a strong AE (FC=2.11, FDR q=0.0008) **(Figures 3B &** 4E, and **Supplementary Table 5).** Our promoter capture HiC data in primary B cells shows an interaction between rs3808619 and the downstream interleukin 7 (*IL7*) gene promoter **(Figure 4F).** *IL7* is a cytokine important for B and T cell development and has been associated with SLE nephritis (19, 20). Taken together, we nominate rs3808619 as a putative causal variant for SLE with potential regulatory action on *IL7*.

Two index SNPs (rs494003 and rs10896045) are reported for the SLE risk haplotype spanning *RNASEH2C* to *OVOL1* on chromosome 11 (21, 22); rs494003 failed to modulate expression in this study **(Figure 5A** and **Supplementary Table 2)**. Multiple SNPs in the region modulated expression, but only two SNPs (rs12293022 and rs10791824 (an hQTL)) produced AEs. Rs12293022 demonstrated significantly increased expression with the alternate T allele (FC=1.62, FDR q=0.036), while the reference A allele of hQTL rs10791824 produced increased expression relative to the alternate G allele (FC=0.61, FDR q=0.038) **(Figures 3B & 5B** and **Supplementary Table 5)**. hQTL rs10791824 is located in intron 2 of *OVOL1,* intersects with *POLR2A* and *CTCF* ChIP-seq peaks, and is in a region of open chromatin in multiple ENCODE datasets **(Supplementary Table 6).** The region containing rs10791824 is shown to interact via our promoter capture HiC data with the promoters of multiple genes besides *OVOL1*, including upstream genes *RELA* and *AP5B1,* and downstream gene *RAB1B* **(Figure 5C).**

**Figure 5.**
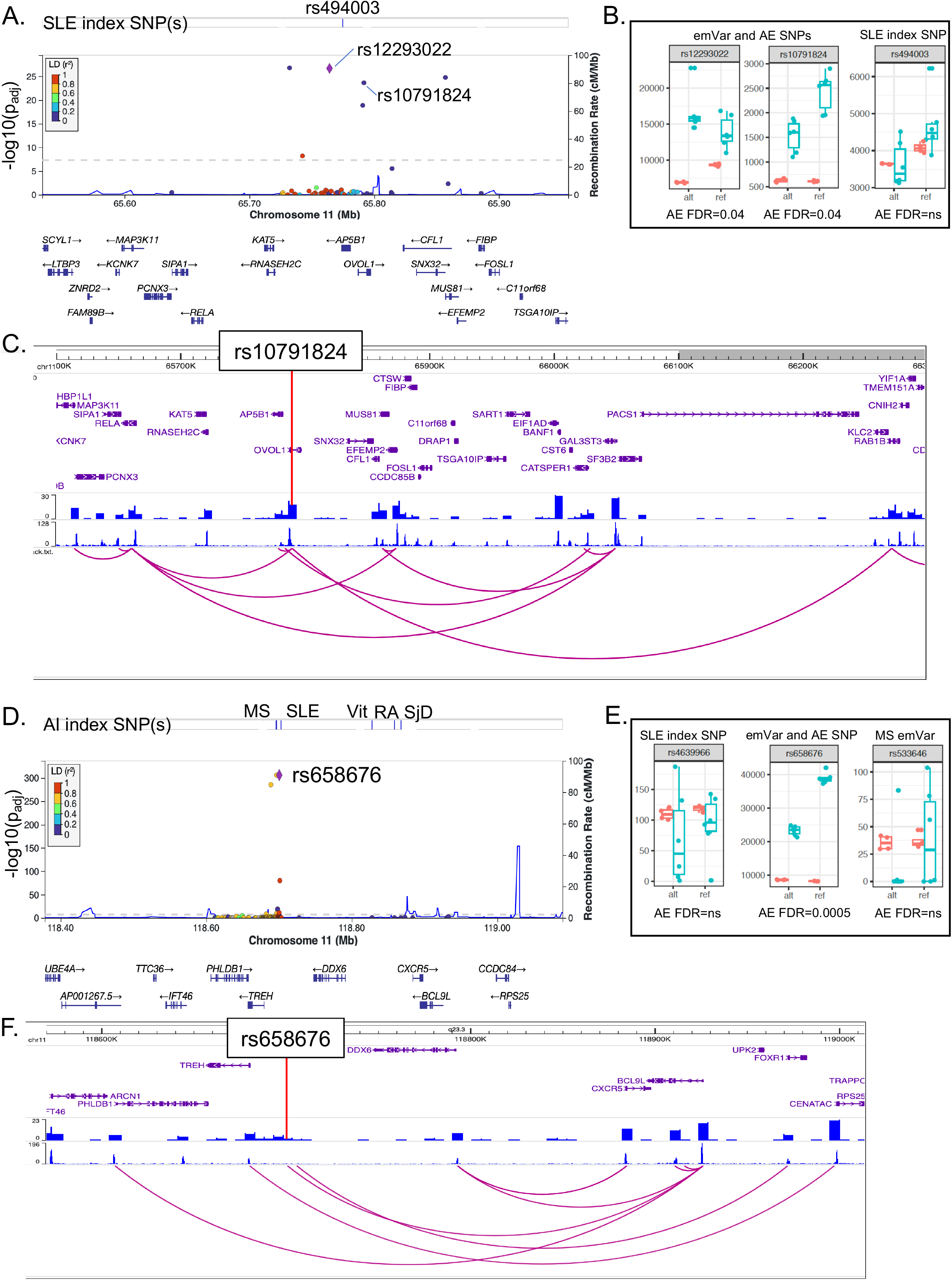
Novel candidate variant identification on the *RNASEH2C-OVOL1* and *PHLDB1-DDX6-CXCR5* SLE risk haplotypes. **A & D.** LocusZoom plots demonstrating emVar effects on the haplotypes of (A) *RNASEH2C-OVOL1* and (D) *PHLDB1-DDX6-CXCR5.* Evaluated index SNPs are indicated. Variants evaluated, their genomic location, and genes in the region are presented on the x-axis. The novel AE variant is represented as a purple diamond. LD values between each variant and the AE variant are colored based on their r^2^ value (see LD key). **B & E.** Box plots of the counts for each allele (alt and ref) in the EBV B (blue) and plasmid control (orange) replicates. FDR q value is indicated. **C & F.** Screenshot from the WashU Epigenome Browser for each haplotype region. Gene positions are provided on the top, followed by density HiC bed tracks, HiC bigwig tracks, and the interactions between gene promoters and the HiC data. The AE variant position is provided.

The region spanning *PHLDB1, DDX6, and CXCR5* on chromosome 11 has been associated with multiple AI diseases. Three SNPs have been associated with SLE (rs4639966, rs480958, and rs4936441) (22, 23). We evaluated 126 variants in this region including rs4639966, as well as index SNPs in the region for MS (rs533646) (24), vitiligo (rs638893) (25), RA (rs10790268) (26, 27), and SjD (rs7119038) (28). None of the tested index SNPs were emVars (**Figure 5D** and **Supplementary Table 2)**. However, rs658676, which is in strong LD with MS index SNP rs533646 (D’=0.962, r^2^=0.888), was a strong emVar and the only tested variant in the region that produced an AE (FC=0.57, FDR q=0.0005 with the reference C allele) **(Figures 3B & 5E** and **Supplementary Table 5).** Our promoter capture HiC data show that the region containing rs658676 interacts with the promoter of B-cell lymphoma 9-like protein (*BCL9L*), a gene just downstream and overlapping a portion of *CXCR5* **(Figure 5F).**

The region on chromosome 16 that spans *CIITA, CLEC16A,* and *SOCS1* is also associated with multiple AI diseases (SLE, celiac disease, MS, T1D, CD, psoriasis, and PBC) (18, 22, 29–46). We evaluated 169 variants in the region including two of the five SLE index SNPs (rs9652601 and rs7200786) and index SNPs for MS (rs2286974, rs7200786, rs12927355), T1D (rs12708716, rs12927355, rs741172, rs2903692), PBC (rs413024, rs1646019), psoriasis (rs367569), and CD (rs423674). Only two index SNPs modulated expression (PBC: rs413024 and CD: rs423674), but neither produced an AE (**Supplementary Tables 2 & 5** and **Supplementary** Figure 3G). Instead, we identified AEs with two other variants in *CLEC16A*, rs1985872 and rs7203793 (D’=0.89, r^2^=0.64), both with the reference alleles displaying significantly higher expression than the alternate alleles (**Supplementary Table 5** and **Supplementary** Figure 4G), prioritizing these variants as candidate causal variants for SLE and multiple other AI diseases.

## Discussion

In this study, we used an MPRA to evaluate the functional role of hQTLs. Our data validate the regulatory potential of 24% (976/4039) of the tested hQTL variants, of which 20.7% demonstrated significant AEs. Only 9.6% (90/942) of variants in strong LD (r^2^>0.8) with hQTLs (hQTL proxies) were AE variants, demonstrating that *a priori* knowledge of hQTLs can facilitate the identification of putative causal variants in the context of strong LD using standard statistical methods. hQTL emVars exhibited significantly stronger effects than eQTL emVars and the other types of variants tested, demonstrating that epigenetic QTLs are more likely to contribute to functional mechanisms than commonly lauded eQTLs. Notably, only 6% of eQTL emVars produced allelic effects, confirming a previous report that only a low proportion of eQTLs overall are likely to be causal (47). Using this approach, we also identified and nominate 17 potential non-HLA causal variants established but incompletely characterized SLE risk haplotypes: *RASGRP3, PTTG1-miR146A, IRF5, BLK, GPR19, IRF8, NEMP2-NAB1, BLK, PKIA-ZC2HC1A, PCNXL3-RNASEH2C, PHLDB1-CXCR5, CABP1, SLC15A4, CIITA-SOCS1, and GRB2.* For several of these regions, we evaluated >150 variants in strong LD, but the MPRA successfully narrowed the important effects to one or two variants per region. We also identified allele dependent regulation for 14 additional published non-HLA AI index SNPs for loci associated with MS, RA, PBC, CD, T1D, KD, UC, GD and vitiligo, prioritizing these variants for future functional studies in B cell lines and B cells.

Our study is the largest MPRA study of SLE to-date, testing 162 SLE index SNPs, 2,477 SNPs on SLE risk haplotypes, and 29,027 additional variants for both functional potential and allelic effect. In 2021, Lu, et al. (10) evaluated 91 known SLE index SNPs and 2,982 variants in strong LD, identifying 51 allelic effects. We expanded upon the study by Lu et al. by testing an additional 81 SLE index SNPs, 2,389 SNPs in LD with SLE index SNPs, and 777 published index SNPs in other AI diseases. Of the 166 variants that were evaluated in both studies, 99 variants (60%) produced concordant regulatory effects: 33 variants were emVars and 66 variants did not modulate expression. Of the 67 variants that exhibited discordant effects, 6 emVars identified in our study and 61 emVars identified by Lu, et al. were not confirmed by the other study. Our study appears to be more conservative in its determinations despite using the same significance and fold change thresholds to call emVars and AE variants. Therefore, we feel confident in the effects that we identify but note that we have likely missed others.

The limitations of this study include the use of EBV B cells because variants that only function in non-B cell types or require specific conditions would not have been detected by our MPRA strategy. Further, because of the minimal promoter used in the MPRA construct, the observed effects were limited largely to inducible rather than suppressive effects on regulatory activity. Lastly, the primary goal of our study was to evaluate hQTLs and was not designed to test every variant in LD with SLE or other AI index SNPs. Therefore, while we successively identified multiple likely causal effects within SLE and AI disease risk haplotypes, there are many others that we have not evaluated. Additionally, selected index SNPs were identified using the GWAS catalog; since risk variants not identified by GWAS or the ImmunoChip arrays (e.g., candidate gene studies or fine mapping studies) are not included in the catalog, they were not considered for this report.

In summary, our study expands our understanding of hQTLs, demonstrating that our tested hQTLs likely regulate aspects of the innate and adaptive immune responses through interactions with IRF TFs, many hQTLs are located in dELS cREs that overlap noncoding RNAs, and a large proportion are likely causal variants for complex trait phenotypes. In addition, we nominate 31 causal variants for SLE and AI diseases. Thus, we uncover important insights into the mechanistic relationships between genotype, epigenetics, gene expression, and SLE and AI disease phenotypes.

## ACKNOWLEDGEMENTS

We would like to thank Dr. Ryan Tewhey for providing the pGL4:23:ΔxbaΔluc and pGL4.23:minP GFP plasmids. We would also like to thank Drs. Ryan Tewhey (The Jackson Laboratory), Leah Kottyan and Xioaming Lu (Cincinnati Children’s Hospital Medical Center), and Kaiyu Jiang (University of Buffalo) for their expert advice regarding different aspects of their MPRA protocols. All sequencing experiments were conducted by the Clinical Genomics Center at the OMRF (https://omrf.org/research-faculty/core-facilities/next-generation-sequencing/). Research reported in this publication was supported by the Presbyterian Health Foundation (Oklahoma City, OK), and the National Institute of Arthritis and Musculoskeletal and Skin Diseases, the National Institute of Allergy and Infectious Diseases, and the National Institute of General Medical Sciences under award numbers R01 AR073606, R01 AI156724, P20 RR020143, P30 AR053483, and P30 GM103510. The content of this publication is solely the responsibility of the authors and does not necessarily represent the official views of the National Institutes of Health or other funding agencies.

## AUTHOR CONTRIBUTIONS

Dr. Gaffney had full access to all of the data in the study and takes responsibility for the integrity of the data and the accuracy of the data analysis.

Study conception and design: Gaffney, Fu, Kelly

Acquisition of data: Fu, Gopalakrishnan, Pasula

Analysis: Kelly, Pelikan

Interpretation of data: Gaffney, Fu, Kelly, Gopalakrishnan, Pelikan, Tessneer

Drafting of article and revising for important intellectual content: Gaffney, Kelly, Tessneer, Fu, Gopalakrishnan, Grundahl, Murphy, Pasula, Pelikan

## ABBREVIATION/DEFINITION LIST

AE: allelic effects; emVars that had both alleles represented and displayed significant differential regulatory effects between the reference and alternate alleles
AI: autoimmune
AP5B1: Adaptor Related Protein Complex 5 Subunit Beta 1
AS: ankylosing spondylitis
Atf3: Activating Transcription Factor 3
BATF: Basic Leucine Zipper ATF-Like Transcription Factor
BCL9L: B-cell Lymphoma 9-like Protein
bHLH: basic helix-loop-helix transcription factor family
bZIP: basic leucine zipper transcription factor family
CD: Crohn’s disease
CIITA: Class II Major Histocompatibility Complex Transactivator
CLEC16A: C-Type Lectin Domain Containing 16A
cREs: cis-regulating elements
CXCR5: C-X-C Motif Chemokine Receptor 5
DDX6: DEAD-Box Helicase 6
dELS: distal enhancer like signature
EBV: Epstein-Barr virus
ELS: enhancer like signature
emVar: expression modulating variant where one or more alleles alter GFP expression
emAlleles: the reference and alternate alleles of an emVar that alter GFP expression
eQTL: expression quantitative trait locus
ETS: erythroblast transformation specific transcription factor family
ETV4: ETS Variant Transcription Factor 4
FC: fold change
FWER: family wise error rate
Fli1: Fli-1 Proto-Oncogene, ETS Transcription Factor
GABPA: GA Binding Protein Transcription Factor Subunit Alpha
GWAS: genome-wide association study
hQTL: histone quantitative trait locus
index SNP: published SNP in GWAS catalog that meets p<5E-08 for association to disease
HLA: human leukocyte antigen
IL7: Interleukin 7
IRF: interferon regulatory factor
JunB: JunB Proto-Oncogene, AP-1 Transcription Factor Subunit
KD: Kawasaki disease
LD: linkage disequilibrium
MAF: minor allele frequency
MFSD6: Major Facilitator Superfamily Domain Containing 6
MPRA: massively parallel reporter assay
MS: multiple sclerosis
NAB1: NGFI-A Binding Protein 1
NEMP2: Nuclear Envelope Integral Membrane Protein 2
OVOL1: Ovo Like Transcriptional Repressor 1
PBC: primary biliary cirrhosis
pELS: promoter enhancer like signature
PHLDB1: Pleckstrin Homology Like Domain Family B Member 1
PKIA: CAMP-Dependent Protein Kinase Inhibitor Alpha
RA: rheumatoid arthritis
RAB1B: RAB1B, Member RAS Oncogene Family
RELA: RELA Proto-Oncogene, NF-KB Subunit
RNASEH2C: Ribonuclease H2 Subunit C
SjD: Sjögren’s disease
SOCS1: Suppressor of Cytokine Signaling 1
SLE: systemic lupus erythematosus
SNP: single nucleotide polymorphism
SS: systemic sclerosis (scleroderma)
T1D: Type 1 diabetes
TF: transcription factor
TSS: transcription start site
UC: ulcerative colitis
ZC2HC1A: Zinc Finger C2HC-Type Containing 1A
ZF: C2H2-zinc factor transcription factor family

