## Supplementary Figures for "Massively Parallel Reporter Assay Confirms Regulatory Potential of hQTLs and Reveals Important Variants in Lupus and Other Autoimmune Diseases"

**Supplementary Material**

**Supplementary Figure 1. MPRA QC and descriptive statistics. A.** Distribution of the number of oligos with > 10 barcodes (n = 66,131; 98.9%) in the aggregated plasmid control libraries. **B.** Distribution of the number of oligos with > 10 barcodes ( n = 61,096, 91.3%) in the aggregated EBV replicate libraries. **C.** Distribution of the number of oligos with > 20 mean counts (n = 65,532; 98%) in the aggregated plasmid control libraries. **D.** Distribution of the number of oligos with > 20 mean counts (n = 64,361, 96.2%) in the aggregated EBV B replicate libraries. **E.** Correlation matrix of oligo counts for each replicate library. **F.** Scatterplot of pairwise comparisons of normalized oligo counts in aggregated plasmid replicates (x-axis) and aggregated EBV replicates (y-axis). Significant emVars are indicated in red.
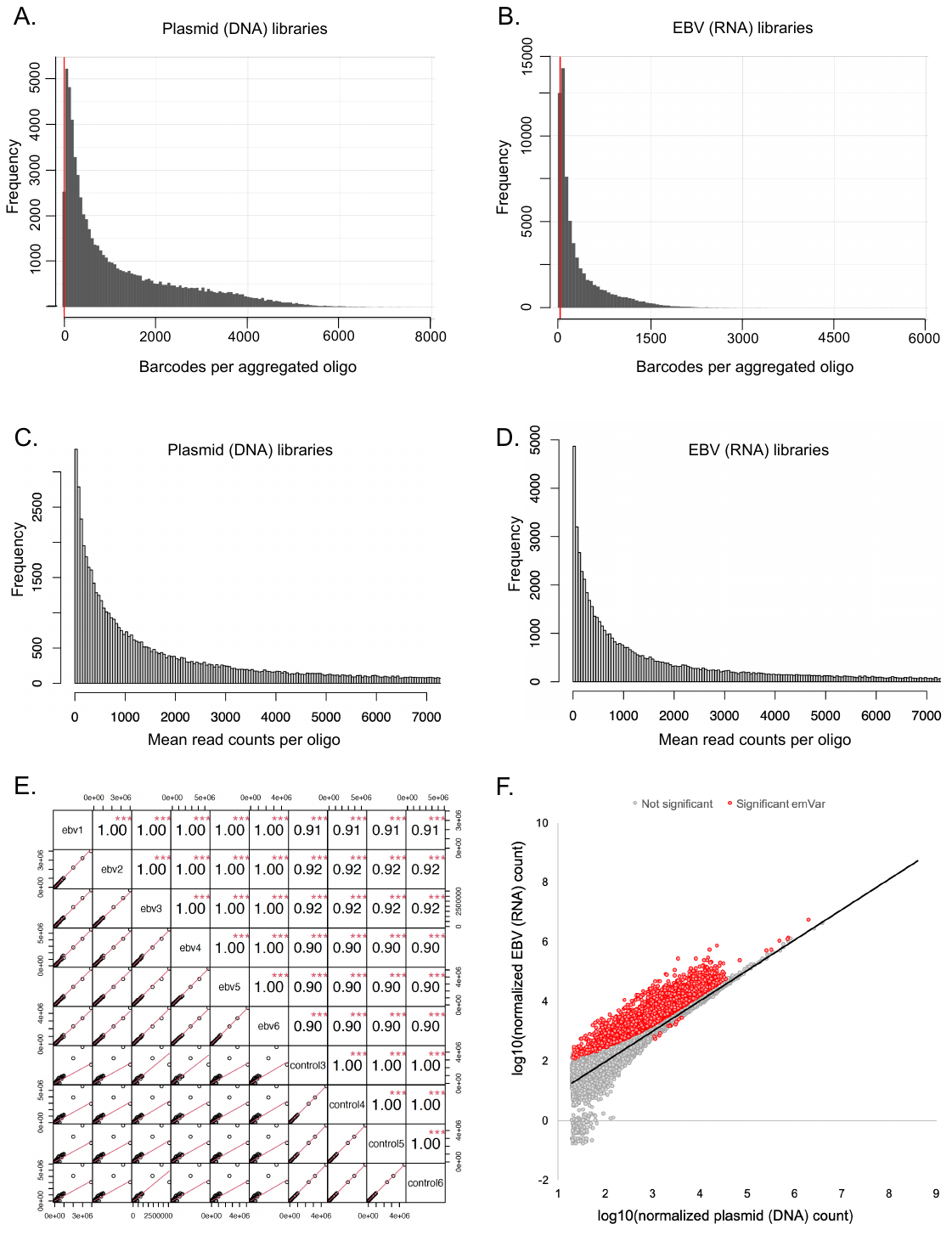

**Supplementary Figure 2. Tested variants located in ENCODE cREs. A.** Proportion (y-axis) of emVars and non-emVars in the different types (x-axis) of cREs. **B.** Proportion (y-axis) of emVars (blue) in cREs by variant type (y-axis). Count is provided above each bar. **C.** Proportion of emVars (gray) in distal enhancer-like signatures by variant type. **D.** Proportion of emVars (green) in proximal enhancer-like signatures by variant type. **E.** Proportion of emVars (purple) in CTCF-only cREs by variant type. **F.** Proportion of emVars (orange) in promoter like signatures by variant type. **G.** Proportion of emVars (yellow) in DNase-H3K4me3 cREs by variant type. Significant differences in proportion of each cRE in hQTLs compared to each other variant type are shown: *<0.05; ** < 0.01, ***<0.001, ****<0.0001.

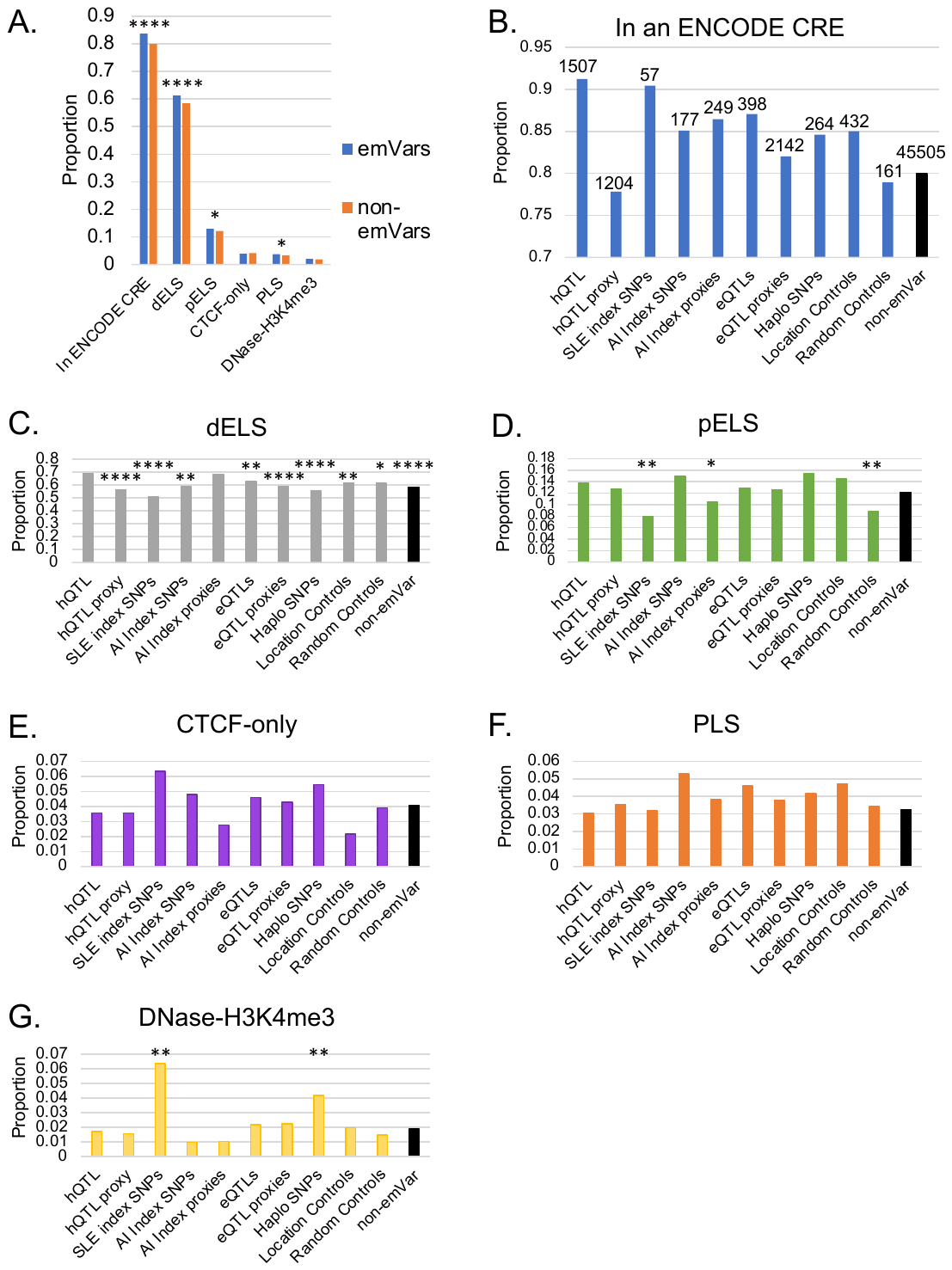

**Supplementary Figure 3. emVar/AE variants on SLE risk haplotypes.** LocusZoom plots demonstrating emVar and allelic effects on SLE risk haplotypes*.* Evaluated index SNPs are presented at the top of graph. Variants evaluated, their genomic location, and genes in the region are plotted on the x-axis. AE variants are represented as a purple diamond. Published SLE index SNP AE variants are circled in green (**A-F**) and novel SLE AEs are circled in red (**G-J**). Variants are colored based on their LD r^2^ values with the circled AE variant (see LD key). The -log_10_(p_adj_) of the emVar score for each variant is plotted on the y-axis.

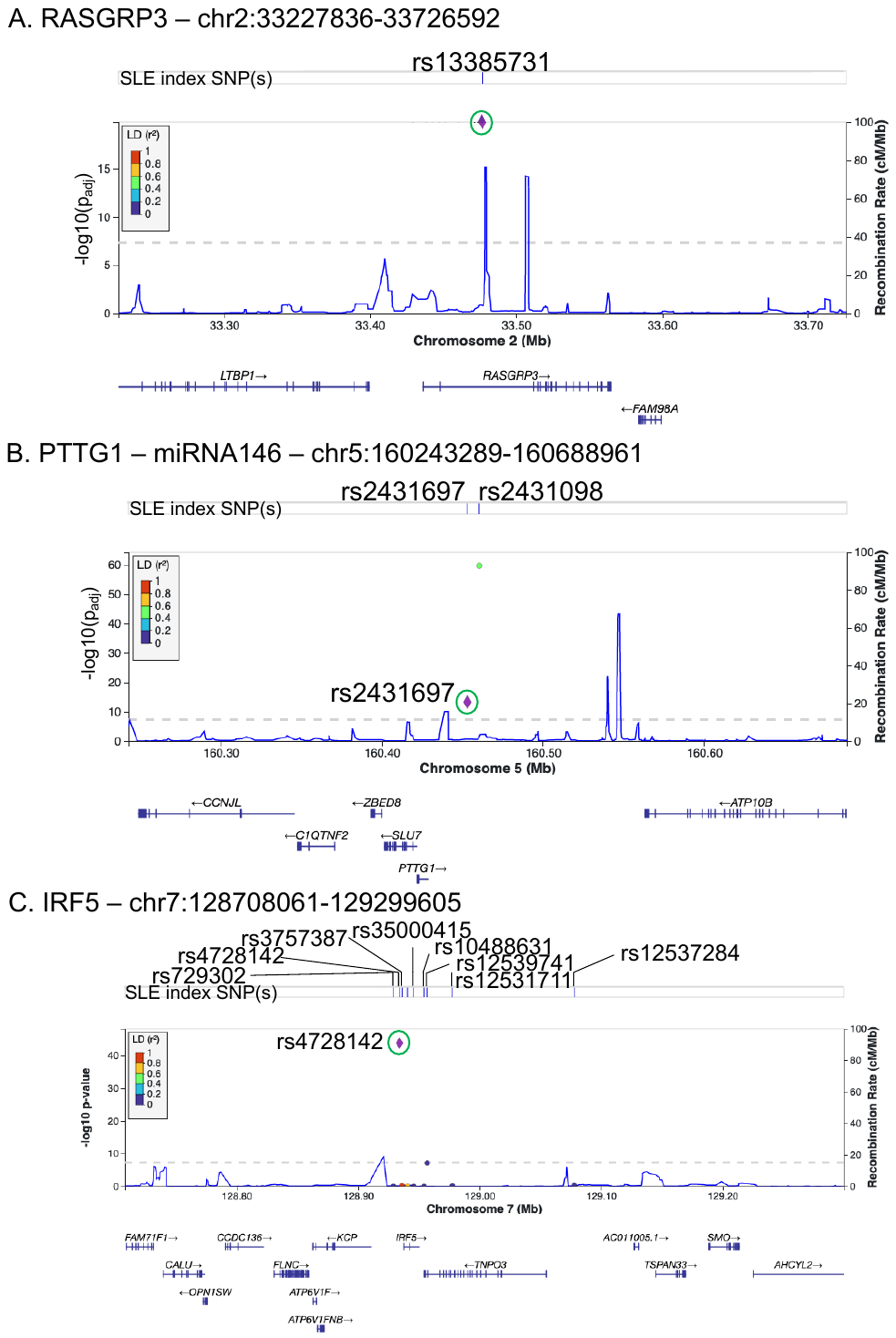

**Supplementary Figure 3 (continued). emVar/AE variants on SLE risk haplotypes.**

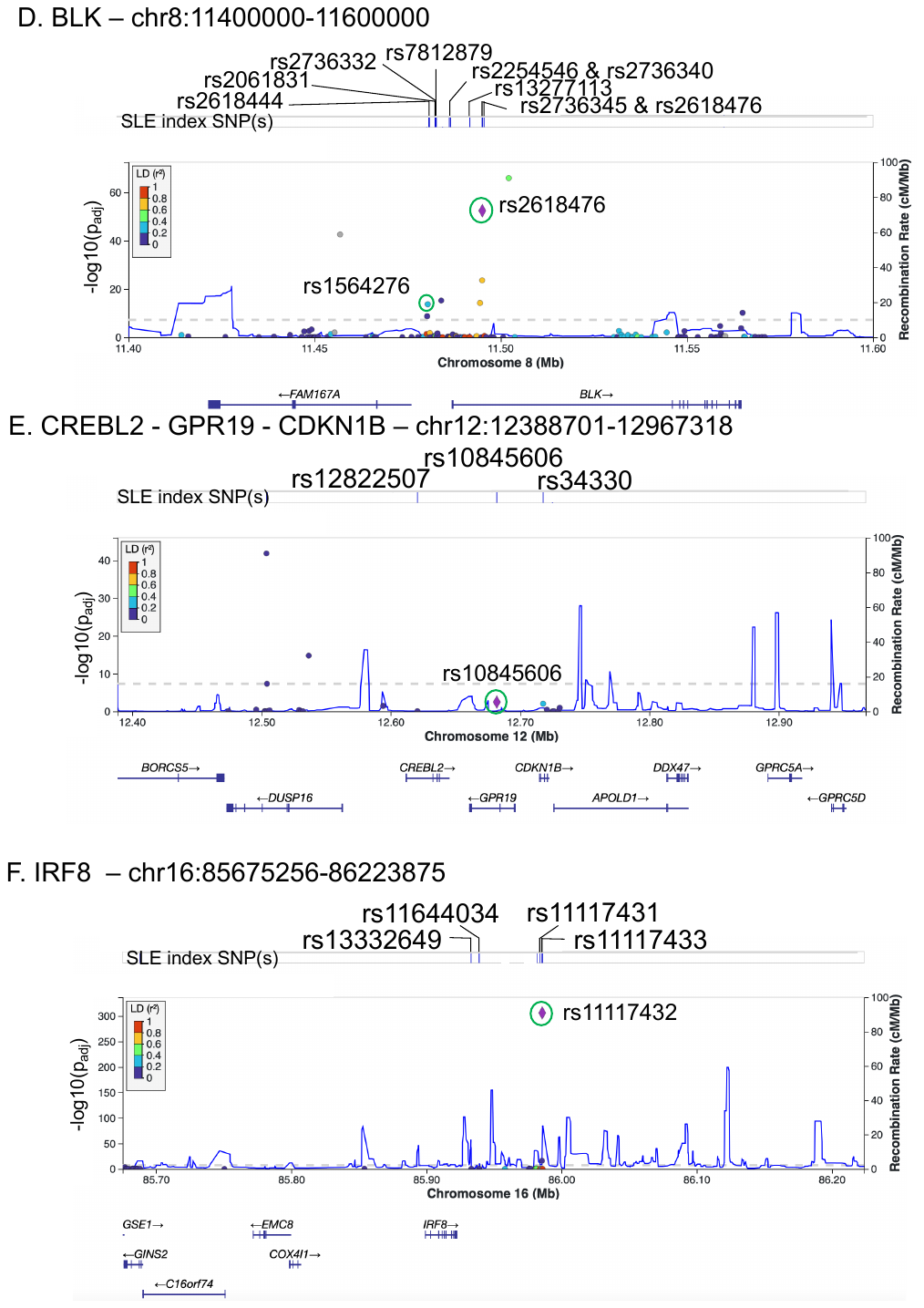

**Supplementary Figure 3 (continued). emVar/AE variants on SLE risk haplotypes.**

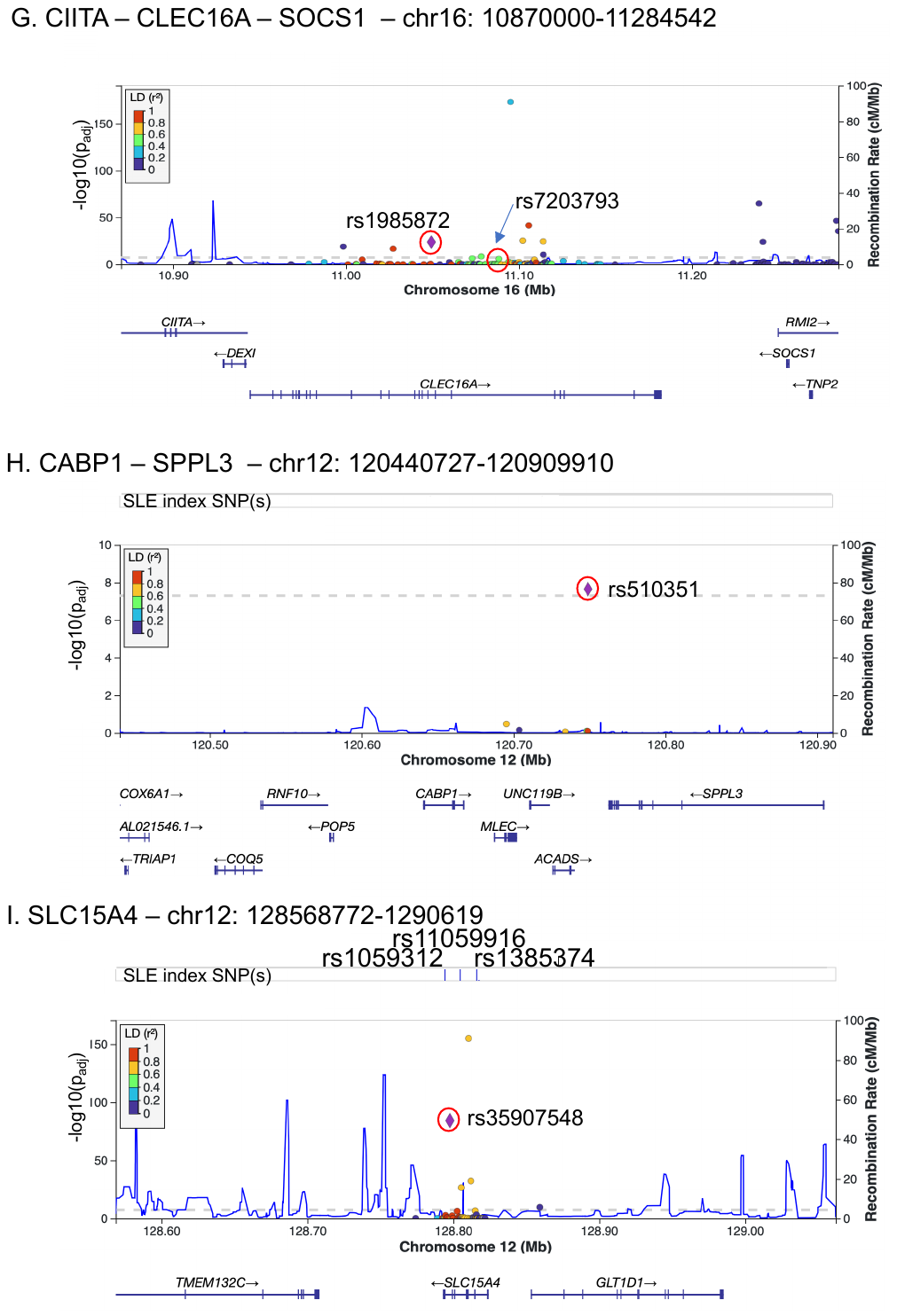

**Supplementary Figure 3 (continued). emVar/AE variants on SLE risk haplotypes.**

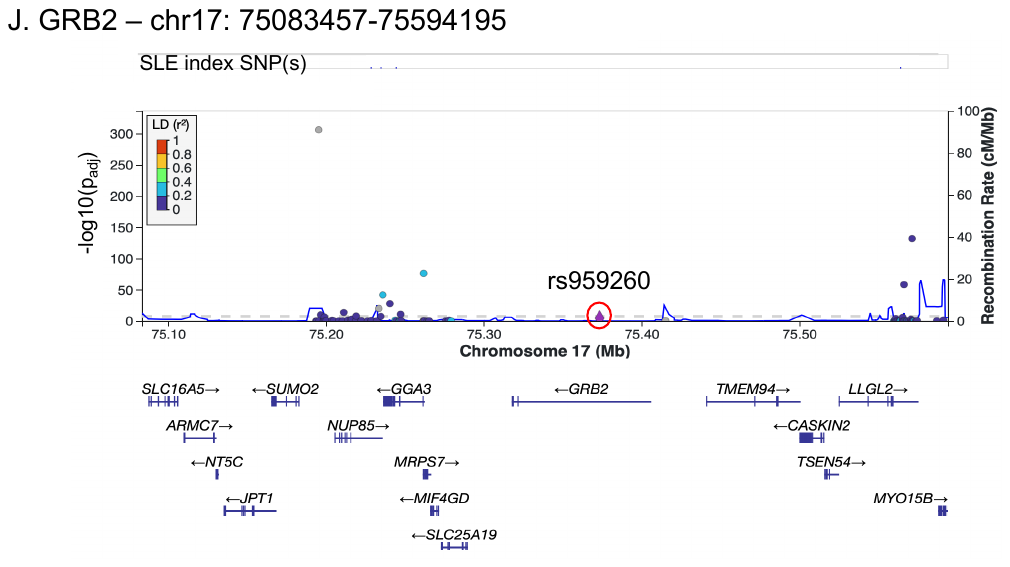

**Supplementary Figure 4. Box plots of SLE allelic effect variants.** Box plots of normalized counts for EBV B replicates (blue) and controls (orange) at each AI allelic variant. Count is plotted on the y-axis and allele (ref/alt) is plotted on the x-axis. The FDR q value and risk gene are provided. EmVars are boxed in blue and AE variants are boxed in orange.

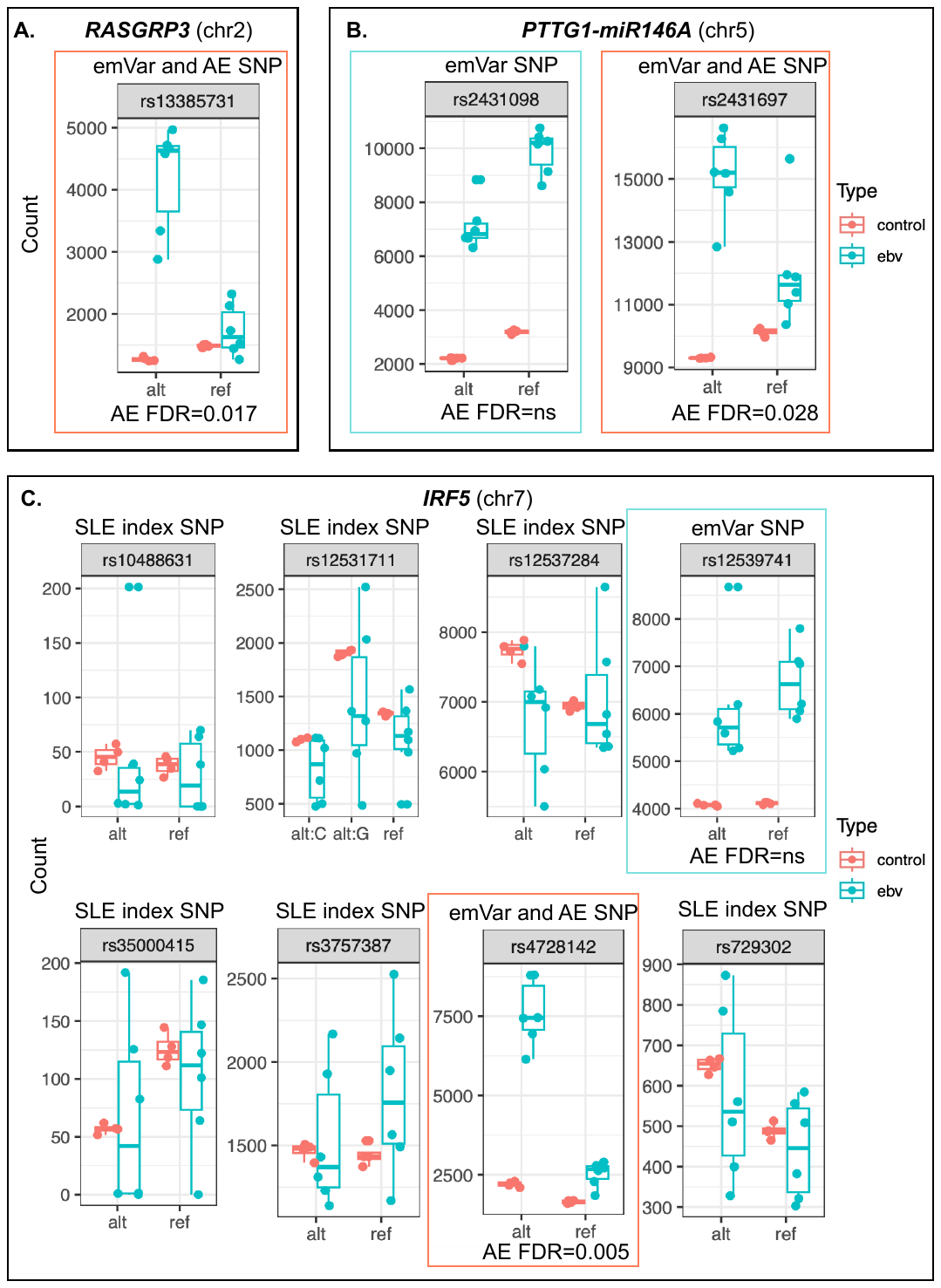

**Supplementary Figure 4 (continued). Box plots of SLE allelic effect variants.**

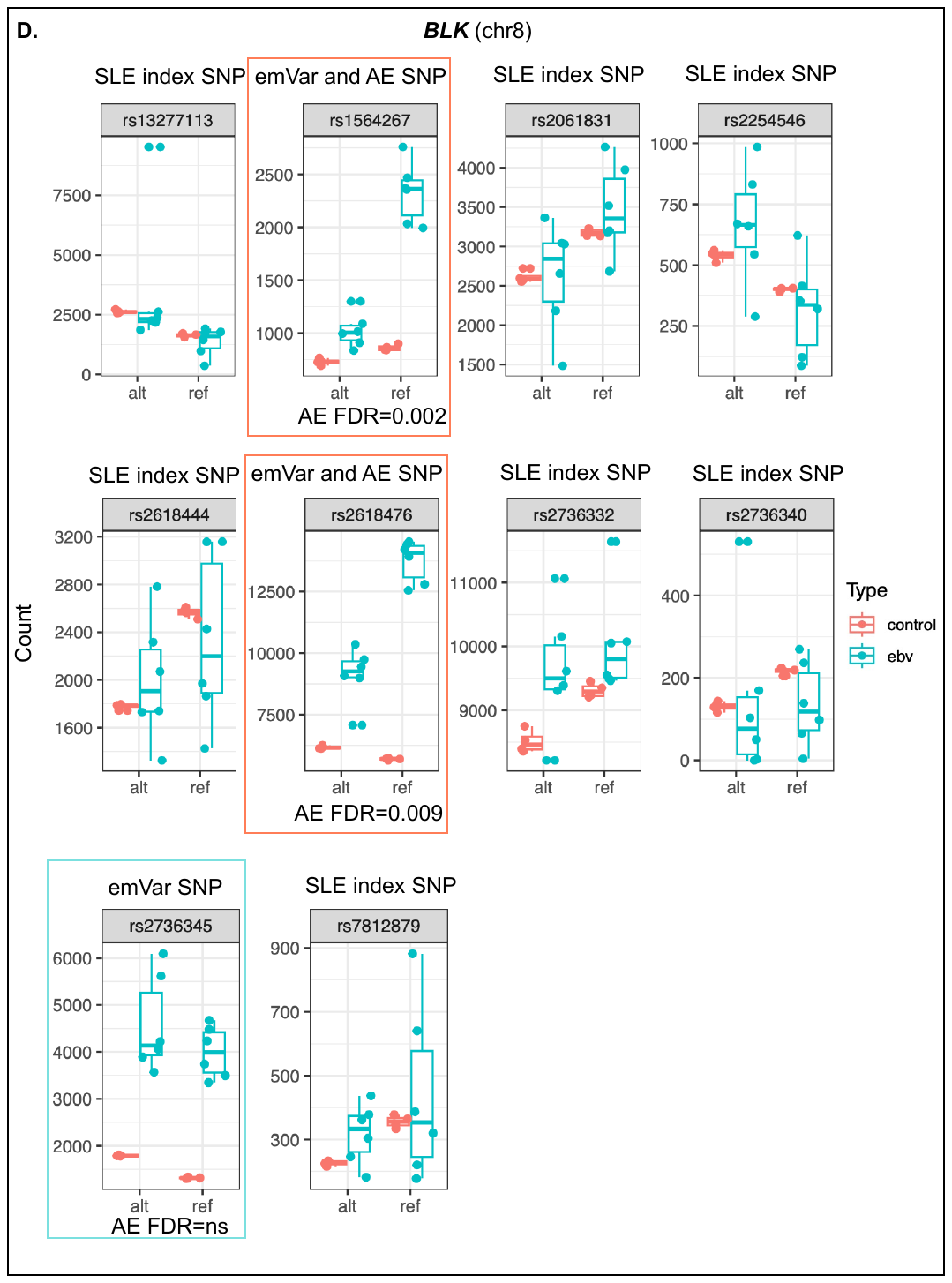

**Supplementary Figure 4 (continued). Box plots of SLE allelic effect variants.**

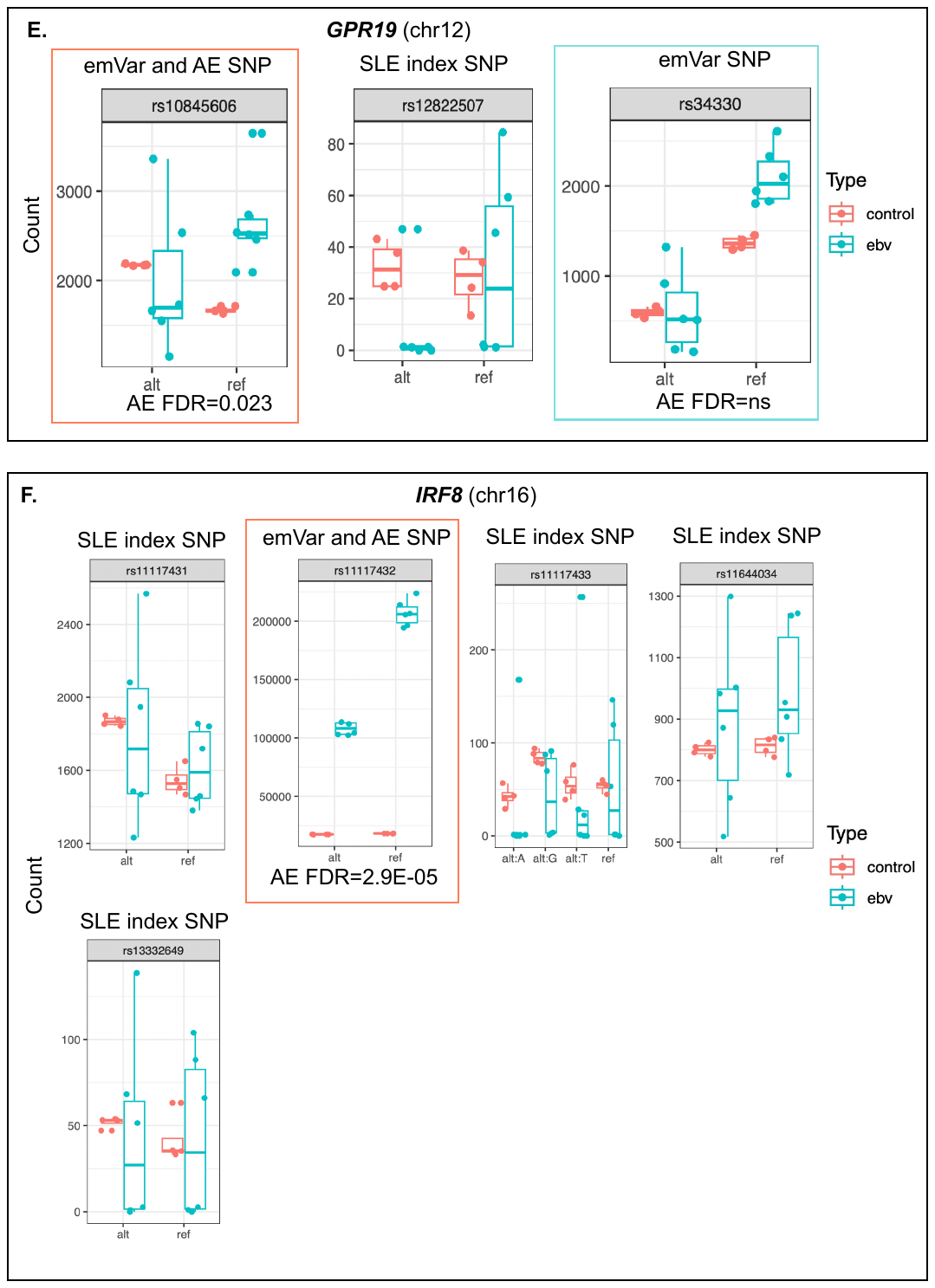

**Supplementary Figure 4 (continued). Box plots of SLE allelic effect variants.**

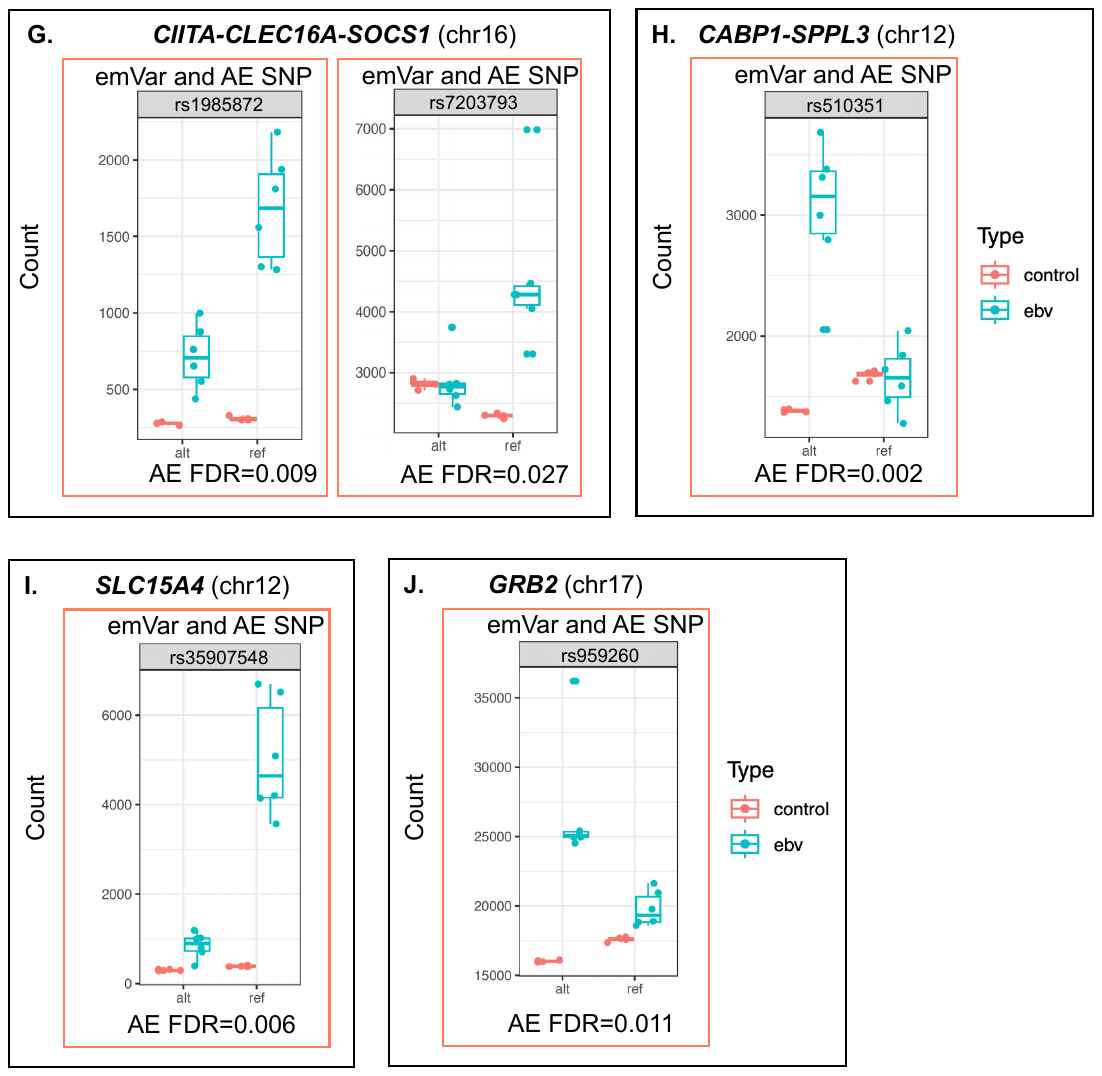

**Supplementary Table 5. Box plots of AI allelic effect variants.** Box plots of normalized counts for EBV B replicates (blue) and controls (orange) at each AI allelic variant. Count is plotted on the y-axis and allele (ref/alt) is plotted on the x-axis. The FDR q value and risk gene are provided. Panels are ordered by rsID.
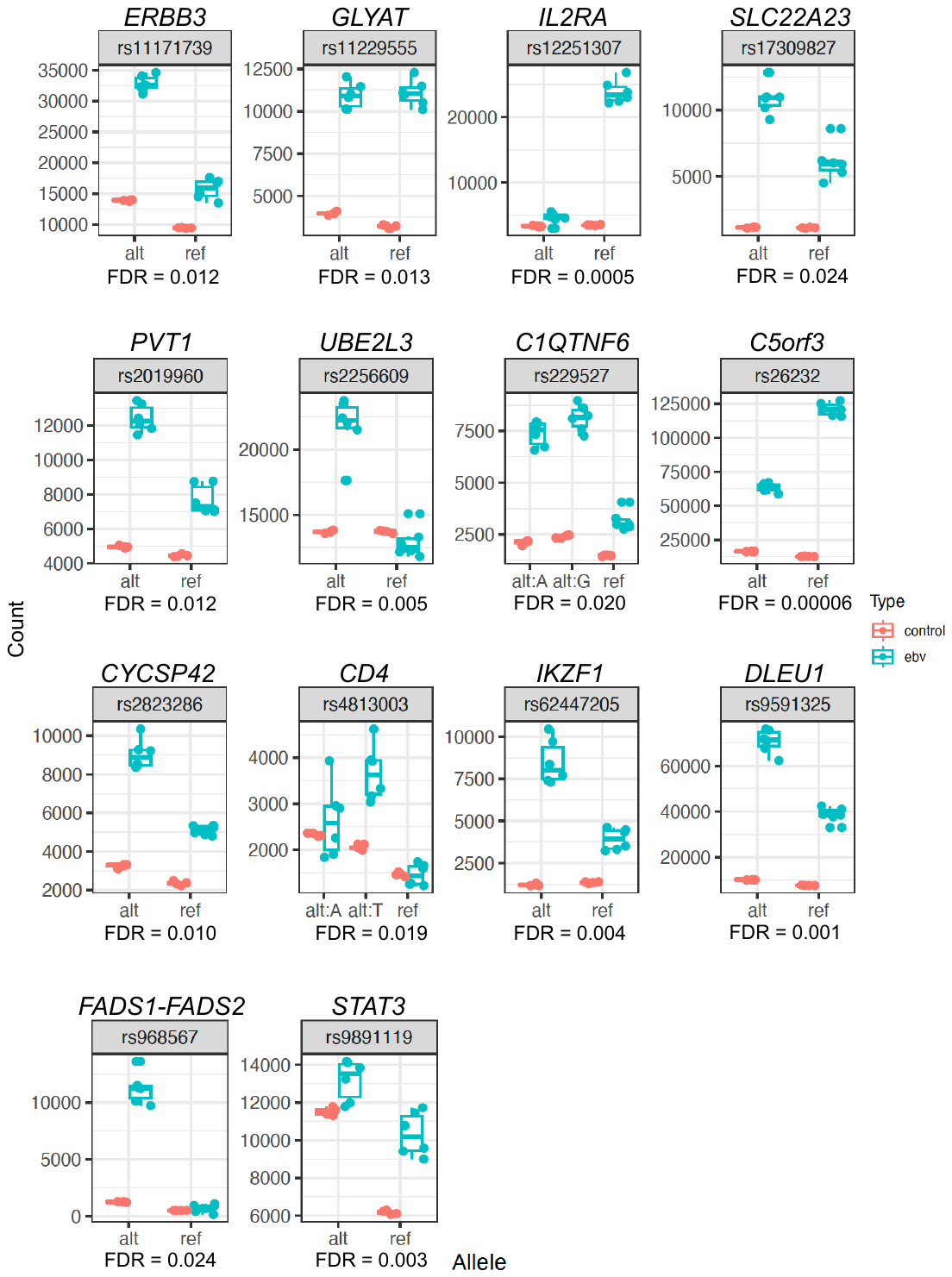
